# Detecting Tumor Specific Antigen-Reactive T cells from Tumor Infiltrating Lymphocytes via Interaction Dependent Fucosyl-biotinylation

**DOI:** 10.1101/2020.03.18.996017

**Authors:** Zilei Liu, Jie P. Li, Mingkuan Chen, Mengyao Wu, Yujie Shi, Wei Li, John R. Teijaro, Peng Wu

**Affiliations:** Department of Molecular Medicine, The Scripps Research Institute, La Jolla, California 92037, United States; State Key Laboratory of Coordination Chemistry, Chemistry and Biomedicine Innovation Center (ChemBIC), School of Chemistry and Chemical Engineering, Nanjing University, Nanjing, China; Department of Oncology, the First Affiliated Hospital of Soochow University, Suzhou, China; Department of Immunology and Microbiology, The Scripps Research Institute, La Jolla, California 92037, United States

## Abstract

Re-activation and clonal expansion of tumor specific antigen (TSA)-reactive T cells are critical to the success of checkpoint blockade and adoptive transfer of tumor-infiltrating lymphocyte (TIL) based therapies. There are no reliable markers to specifically identify the repertoire of TSA-reactive T cells due to their heterogeneous composition. Here we introduce FucoID as a general platform to detect endogenous antigen-specific T cells and study their biology. Through this interaction dependent labeling approach, TSA-reactive T cells can be detected and separated from bystander T cells in primary tumor digests based on their cell-surface enzymatic fucosyl-biotinylation. Compared to bystander TILs, TSA-reactive TILs possess a distinct TCR repertoire and unique gene features. Though exhibiting a dysfunctional phenotype, this subset of TILs possesses substantial capabilities of proliferation and tumor specific killing. FucoID features genetic manipulation-free procedures and a quick turnover cycle, and therefore should have the potential of accelerating the pace of personalized cancer treatment.

**Highlights:** Interaction dependent fucosylation enables the detection and isolation of *bona fide* intratumoral tumor specific antigen-reactive T cells

Tumor specific antigen-reactive CD8^+^ T cells possess capabilities to be expanded and adoptively transferred for tumor control

Tumor specific antigen-reactive CD8^+^ T cells feature oligoclonal expansion and upregulate genes for the steroid biosynthesis and metabolic process

Intratumoral bystander CD8^+^ T cells can be separated into two groups based on PD-1 expression that feature distinct gene modules

## Introduction

In the past decade, the development of immune checkpoint inhibitors and adoptive cell transfer (ACT)-based therapies has revolutionized cancer treatment (Chowdhury et al., 2018; Guedan et al., 2018; Rosenberg and Restifo, 2015; Sharma and Allison, 2015). The success, however, is limited to a relatively small subset of patients and cancer types (Galon and Bruni, 2020; Sanmamed and Chen, 2018; Yamamoto et al., 2019). Successful anti-tumor immune responses following immunotherapy are believed to require re-activation and clonal expansion of tumor specific antigen (TSA)-reactive T cells present in the tumor microenvironment (Gubin et al., 2014; McGranahan et al., 2016; Ott et al., 2017; Rizvi et al., 2015; Sahin et al., 2017; Schumacher and Schreiber, 2015). The unpredictability of a patient’s response to immunotherapy is partially attributed to the heterogeneity of the tumor immune microenvironment and phenotypic profiles of TILs within individual tumors. (Chevrier et al., 2017; Lavin et al., 2017; Rizvi et al., 2015; Stevanović et al., 2017) Previous studies revealed that TILs consist of not only those T cells specific for TSAs, but also the ones that recognize epitopes unrelated to the tumor, such as virus specific antigens, known as bystander T cells. (Scheper et al., 2019; Simoni et al., 2018) Therefore, identifying TSA-reactive T cells from cancer patients has critical implications for prediction and therapeutic applications. Although tumor-reactive TIL candidates can be roughly enriched from tumor digests through the expression of cell-surface markers (e.g. PD-1) (Gros et al., 2016; 2014a; Yossef et al., 2018) (Gros et al., 2016; 2014b; Yossef et al., 2018), bystander T cells expressing such makers also exist in the tumor microenvironment (Duhen et al., 2018; Sade-Feldman et al., 2018; Scheper et al., 2019; Simoni et al., 2018).

The precise therapeutic effect of immunotherapies that boost immunity of endogenous T cells is governed by the T cells’ capability to recognize TSAs (Schumacher and Schreiber, 2015). At the molecular level, this is determined by the interaction of unique T cell receptor (TCRs) with cognate peptide–major histocompatibility complexes (pMHCs) (Dembić et al., 1986). Designed based on this knowledge, peptide–MHC (pMHC) multimers are widely used to profile TCR specificity of a known antigen and to identify TILs specific to a particular TSA (Cohen et al., 2015; Echchakir et al., 2002; Glanville et al., 2017; Newell et al., 2013; Robbins et al., 2013; Yamamoto et al., 2019). To identify TSAs for the pMHC approach, however, hinges on reverse immunology, in which whole exome sequencing (WES) is performed on tumor cells to uncover nonsynonymous mutations (Robbins et al., 2013; Schumacher and Schreiber, 2015). *In silico* tools are then used to generate peptides harboring epitopes encoded by these identified nonsynonymous mutations. Such peptides are either left unfiltered, filtered through the use of prediction algorithms for MHC-binding, or used as guides for identifying MHC-associated TSAs by combining with mass spectrometry analysis (Abelin et al., 2017; Yamamoto et al., 2019). Such methods have enabled the identification of TSAs and TSAs-reactive TCRs for melanoma patients and a small number of patients with epithelial cancers (Stevanović et al., 2017; Tran et al., 2016; Zacharakis et al., 2018). Despite these acclaimed successes, a large portion of identified TSA candidates is not immunogenic because available computational tools cannot accurately predict T-cell reactivity (Editorial, 2017). Requiring deep sequencing, bioinformatics analysis and machine learning, these methods heavily rely on expertise in computational biology, and, as a result, turnover times are long (Arnaud et al., 2020; Editorial, 2017; Lee et al., 2018; Liu and Mardis, 2017). Therefore, to accelerate the pace of cancer immunotherapy a computation-free method that enables quick TSA-reactive T cell discovery and easy adoptability would be a welcome advance.

Here we report the development of a method for the identification of antigen-specific T cells based on an interaction dependent fucosyl-biotinylation and the use of this method to isolate endogenous TSA-reactive T cells from tumor digests without the previous knowledge of the TSA identities. In this approach, which we termed FucoID, tumor-lysate primed DCs presenting TSAs, serving as “living” tetramers, are equipped with an enzyme that induces proximity-based transfer of fucosylated biotin (Fuc-Bio) tags to the surface of T cells that interact with the DCs. We demonstrate that the tagged CD8^+^ T cells are bona fide TSA-reactive T cells and based on their cell-surface fucosyl-biotinylation they can be readily separated from bystander T cells. We discover that nearly all TSA-reactive CD8^+^ T cells co-express PD-1, whereas bystander T cells consist of PD-1^+^ and PD-1^−^ subsets. In comparison to bystander TILs, TSA reactive TILs possess a distinct TCR repertoire and unique gene features that are characterized by a dysfunction/activation transcript profile and genes upregulated in steroid biosynthesis and related metabolic pathways. By contrast, genes associated with antiviral defense mechanisms are enriched in bystander T cells. Despite exhibiting a dysfunctional phenotype, the TSA reactive TILs (i.e. PD-1^+^Bio^+^) possess substantial capabilities to proliferation. Upon expansion, this subset of TILs leads to significantly more efficient tumor killing than the expanded, entire PD-1^+^ TIL population both *in vitro* and *in vivo*.

Compared to the techniques that rely on bioinformatics-assisted TSA identification, FucoID features much simpler procedures and a quicker turnover cycle. Importantly, this method opens up an avenue for the characterization and manipulation of the entire repertoire of endogenous, TSA-reactive TILs, and is generally applicable to a number of murine tumor models with detectable T-cell infiltration, including those of MC38 colon cancer (a valid model for hypermutated colorectal cancer) (Efremova et al., 2018) and E0771 triple negative breast cancer (TNBC) whose human counterparts only showed a 12-19% objective response rate to checkpoint inhibitors (Crosby et al., 2018; Hendrickx et al., 2017), and has a high potential to be tested in a clinical setting.

## Results

### Install *H. pylori* α1-3-Fucosyltransferase (FT) onto the cell surface for probing cell-cell interactions

Although a number of enzyme-based proximity-labeling systems have been developed to profile protein-protein interactions(Branon et al., 2018; Kim and Roux, 2016; Liu et al., 2018; Long et al., 2016; Slavoff et al., 2011), very few can be applied to probe cell-cell interactions(Ge et al., 2019; Liu et al., 2018; Pasqual et al., 2018). To the best of our knowledge, all of these approaches rely on genetic manipulations. To design an enzymatic approach for probing cell-cell interactions of primary cells including those of the human origin requires an enzyme that can be installed onto the cell surface of bait cells without genetic engineering. For general applicability, the acceptor substrate of this enzyme should be naturally present on the cell surface of most cell types. The challenge is to identify an enzyme to achieve intercellular labeling with high sensitivity when two cells interact and low background in the absence of an interaction.

Previously, we developed a chemoenzymatic method that enables us to conjugate proteins onto the cell surface (Li et al., 2018). This method relies on α1-3-fucosyltransferase (FT), a glycosyltransferase possessing remarkable donor substrate tolerance. It enables rapid and quantitative transfer of proteins conjugated to the enzyme’s natural donor substrate GDP-fucose (GDP-Fuc) to LacNAc (and/or α2,3-sialyl LacNAc), a common building block of the glycocalyx, on the surface of live cells. Because LacNAc and sialyl LacNAc are abundantly expressed by most cell types, including dendritic cells (DCs) and T cells, this method serves as a general approach for engineering the cell-surface landscape of immune cells. Based on this technique, we devised a genetic-engineering free strategy to probe cell-cell interactions using the membrane anchored FT that is introduced to the cell surface via the chemoenzymatic approach (**Fig. 1A and B**). Due to its high *K*_*m*_ (1.3 mM) and high *k*_*cat*_ (442 min^-1^) (Soriano del Amo et al., 2010; Zheng et al., 2011), the membrane anchored FT is ideal to enable proximity-dependent labeling of prey cells that interact with bait cells harboring the enzyme (**Fig. 1A**). In the absence of an interaction, the concentration of LacNAc (sLacNAc) acceptors in the proximity of FT is far below the FT-LacNAc *K*_*m*_. Under such circumstances, the bimolecular reaction rate is governed by *k*_*cat*_/*K*_*m*_. By having a high *K*_*m*_, the background labeling is minimized. When two cells interact, the local concentration of LacNAc (sLacNAc) in the vicinity of FT is high such that the pseudo-zero-order reaction rate is determined by *k*_*cat*_. By having a high *k*_*cat*_, the labeling signal in the presence of a *bona fide* cell-cell interaction is maximized.

**Figure 1.**
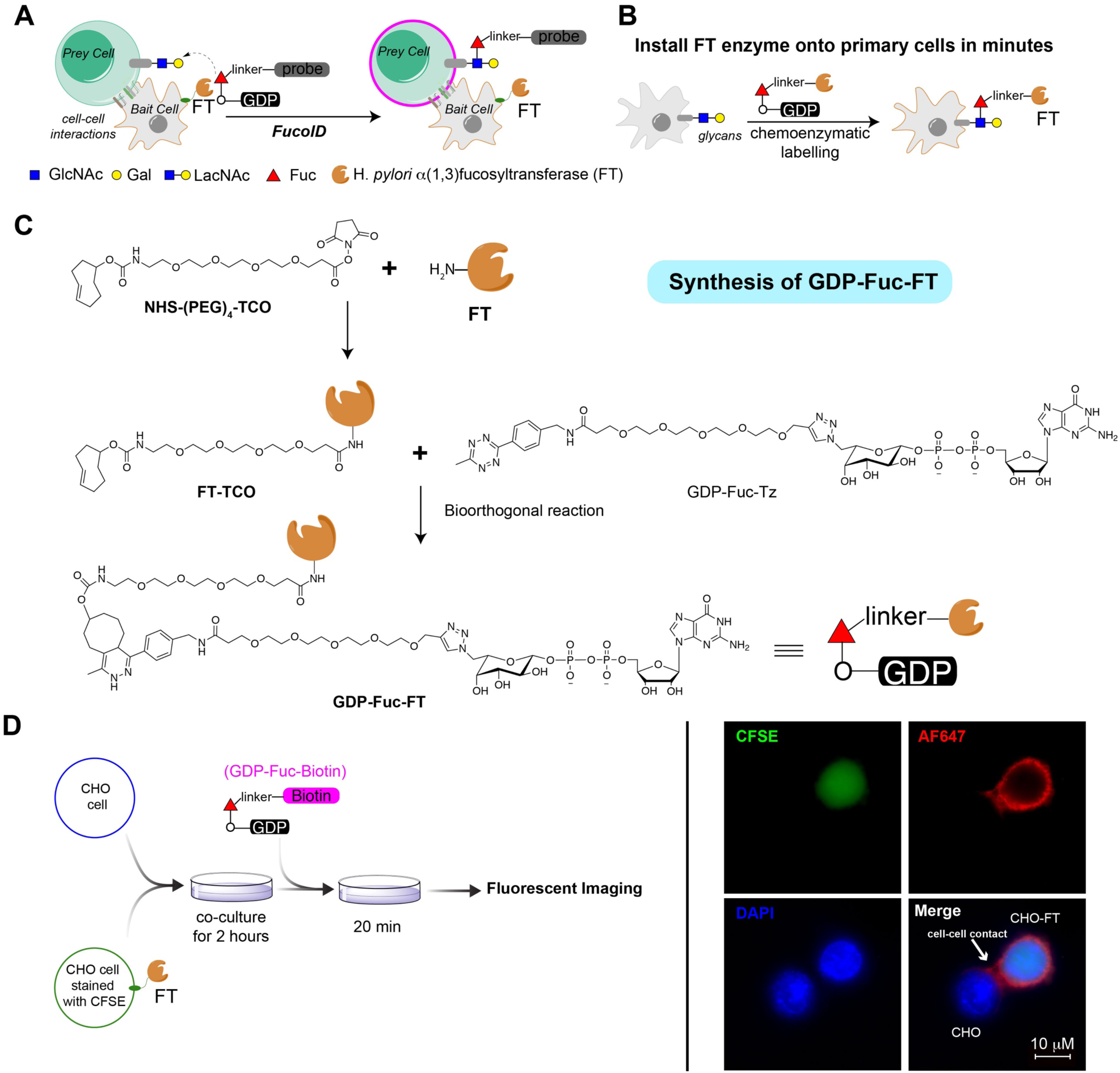
Illustration of probing cell-cell interactions via interaction dependent fucosylation (FucoID) (**A**) Schematic representation of FucoID for the labeling of cell-cell interactions. (**B**) Conjugation of *H. pylori* α(1,3)fucosyltransferase (FT) onto the cell surface via the FT-mediated chemoenzymatic glycan labelling. (**C**) Synthesis of GDP-Fuc-FT. (**D**) Experimental design and the representative fluorescent microscopy image of the proximity labelling mediated by FT functionalized CHO cells (CHO-FT). CHO-FT cells stained with CFSE were mixed (cell ratio 1:5) with unfunctionalized CHO cells and cultured for 2 hours, followed by the addition of GDP-Fuc-Biotin (50 µM). Cells were stained with DAPI and streptavidin-APC for fluorescent microscopy imaging.

As shown in **Fig. 1B**, when cells were incubated with GDP-fucose-conjugated FT (GDP-Fuc-FT) (**Fig. 1C)**, the donor substrate modified enzyme served as a self-catalyst to transfer Fuc-FT onto the cell surface LacNAcylated glycans (**Fig. S2 and 3**). Through this method, primary mouse CD8^+^ T cells and bone marrow derived DCs, human lymphocytes and DCs derived from human PBMC were successfully conjugated with FT and the robust modifications were achieved within 20 min (**Fig. S3A**). Importantly, the installed FT remained on the cell surface for approximately 10 hours (**Fig. S3C**) and did not affect the cell viability (**Fig. S3D**) and functions, e.g. DC-mediated CD8^+^ T cell priming (**Fig. S4**).

Next, we assessed if FT-functionalized cells could mediate intercellular labeling by transferring Fuc-Bio from exogenously added GDP-fucose-biotin (GDP-Fuc-Bio, structure shown in **Fig. S1**) to interacting cells. We incubated FT-functionalized wild-type CHO cells (CHO-FT) with adherent CHO cells to form of cell-cell contacts. Upon the addition of GDP-Fuc-Bio, fluorescence microscopy imaging revealed that unfunctionalized CHO cells not in contact with CHO-FT were not labelled, but the CHO cells in contact with CHO-FT were strongly labelled at the cell-cell contacting interface (**Fig. 1D; Fig. S5**). Not surprisingly, CHO-FT cells were also robustly labelled on the membrane via self-fucosyl-biotinylation **(Fig. 1D**).

### Probe DC-T cell interactions via FucoID

To determine if FucoID could be applied to probe antigen specific DC-T cell interactions, we first used a model system to examine FT-modified immature DCs (iDCs) for the selective labeling of CD8^+^ T cells from OT-I transgenic mice that express a transgenic T-cell receptor (TCR) specific for the SIINFEKL peptide (OVA_257-264_) of chicken ovalbumin presented on MHC-I (**Fig. 2A**). FT-modified iDCs (CD45.1^+/+^) were primed with OVA_257-264_ before co-culturing with naïve OT-I CD8^+^ T cells (CD45.2^+/+^), followed by the addition of GDP-Fuc-Bio to enable the labeling. Robust Fuc-Bio labeling was found on the interacting OT-I CD8^+^ T cells with a signal-to-background ratio of 36% versus 1% (**Fig. 2B**). Here, the background is defined as the signal produced on OT-I CD8^+^ cells by incubating OVA_257-264_-primed iDCs (without membrane-anchored FT) with naïve OT-I CD8^+^ for the same period of time. By contrast, the iDC-FT loaded with the lymphocytic choriomeningitis virus (LCMV) GP_33-41_ peptide only induced the background level labeling of OT-I CD8^+^ T cells, indicating the labeling is highly specific (**Fig. 2B**). As the iDC-FT to T cell ratio increased, the intensity of the FT-mediated intercellular labelling increased accordingly, this was accompanied by the concomitant upregulation of the T-cell early activation marker CD69 (**Fig. 2C**).

**Figure 2.**
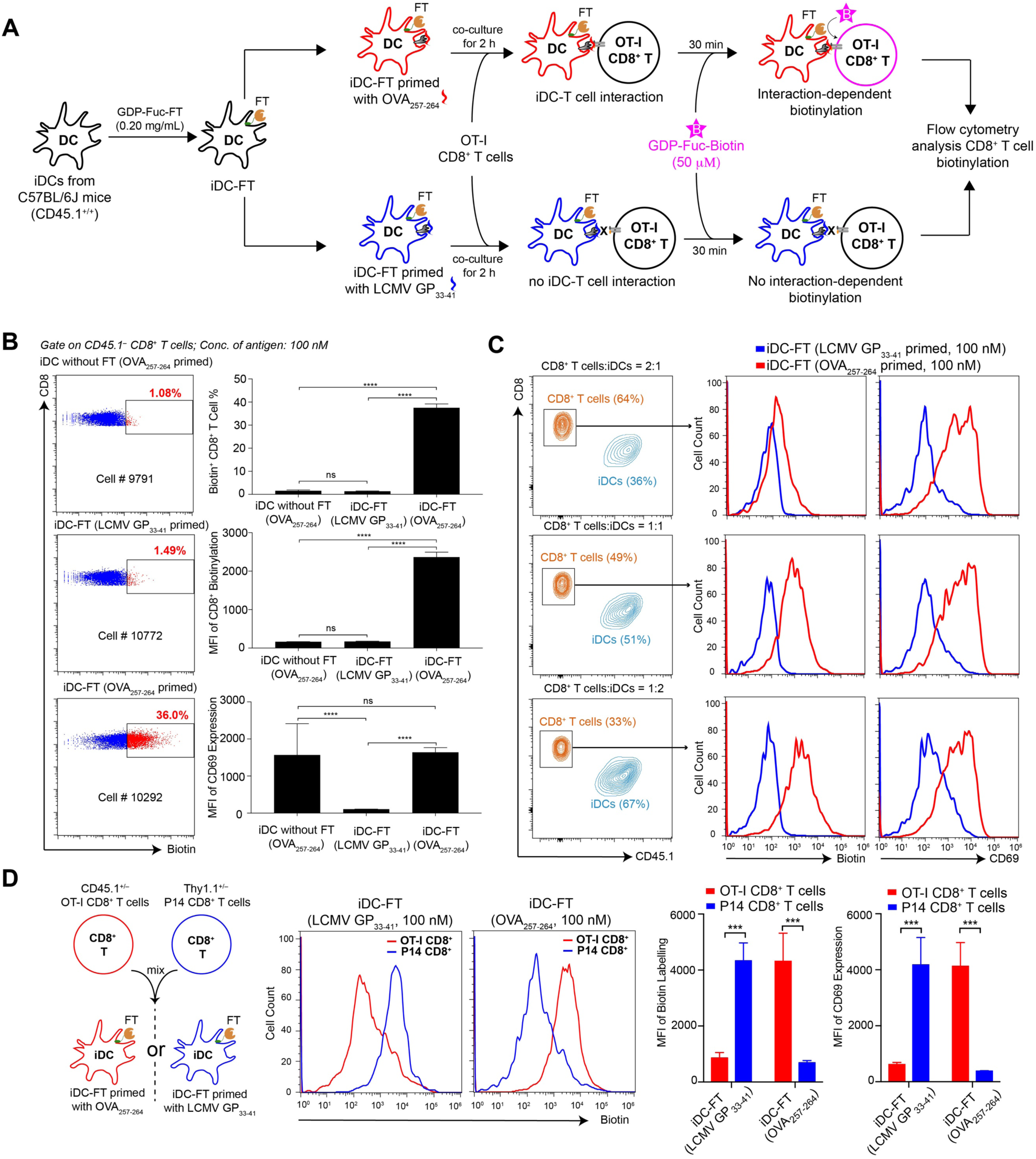
Demonstration of the specificity of FucoID in probing DC-T cell interactions. **(A)** Workflow of analyzing the FucoID enabled proximity labeling signal of naïve OT-I CD8^+^ T cells under the interaction with LCMV GP_33-41_ or OVA_257-264_ primed iDCs. **(B)** Flow cytometric analysis showing antigen specific Fuc-biotinylation of CD8^+^ T cells under different iDC/T ratio of 1:1. **(C)** Flow cytometric analysis showing antigen specific Fuc-biotinylation of OT-I CD8^+^ T cells at different iDC/T cell ratios. Representative figures from three independent experiments. **(D)** Flow cytometric analysis of antigen-specific Fuc-biotinylation in the mixture of OT-I and P14 CD8^+^ T cells when incubated with iDC-FT primed with LCMV GP_33-41_ or OVA_257-264_. In all figures, mean ± SD (n=3); ns, P > 0.05; *P < 0.05; **P < 0.01; ***P < 0.001; ****P < 0.0001; one-way ANOVA followed by Tukey’s multiple comparisons test; two-way ANOVA followed by Sidak’s multiple comparisons test.

To further assess the sensitivity and specificity of FucoID in a more challenging situation, we mixed naïve OT-I (CD45.1^+/–^) and P14 CD8^+^ T cells (Thy1.1^+/–^) that recognize LCMV GP_33-41_ presented by MHC-I and co-cultured the mixture with iDC-FT pulsed with OVA_257-264_ or LCMV GP_33-41_, respectively at the ratio of 1:1:1 (**Fig. 2D**). After co-culturing for 2 hours, GDP-Fuc-Biotin was added. As illustrated in **Fig. 2D**, OT-I and P14 CD8^+^ T cells were selectively labeled by OVA_257-264_ and LCMV GP_33-41_ primed iDCs-FT, respectively.

Next, we sought to determine if antigen specific labeling via FucoID could be achieved in cell mixtures of natural components and complexity. To this end, iDCs-FT primed by OVA_257-264_- and LCMV GP_33-41_, respectively, were co-cultured with OT-I splenocytes. As expected, significant OVA-specific fucosyl-biotinylation were detected (44 % versus 0.6%) (**Fig. 3A and B**). Subsequently, we optimized the labeling condition using the iDC–OT-I splenocyte co-culturing system, finding that the optimal labeling was achieved within 2 hours using iDC modified with 0.2 mg/mL GDP-Fuc-FT (**Fig. S6A and B**). Under this condition, the antigen specific labeling was positively correlated with the amount of OVA_257-264_ for iDC priming (**Fig. S6C**).

**Figure 3.**
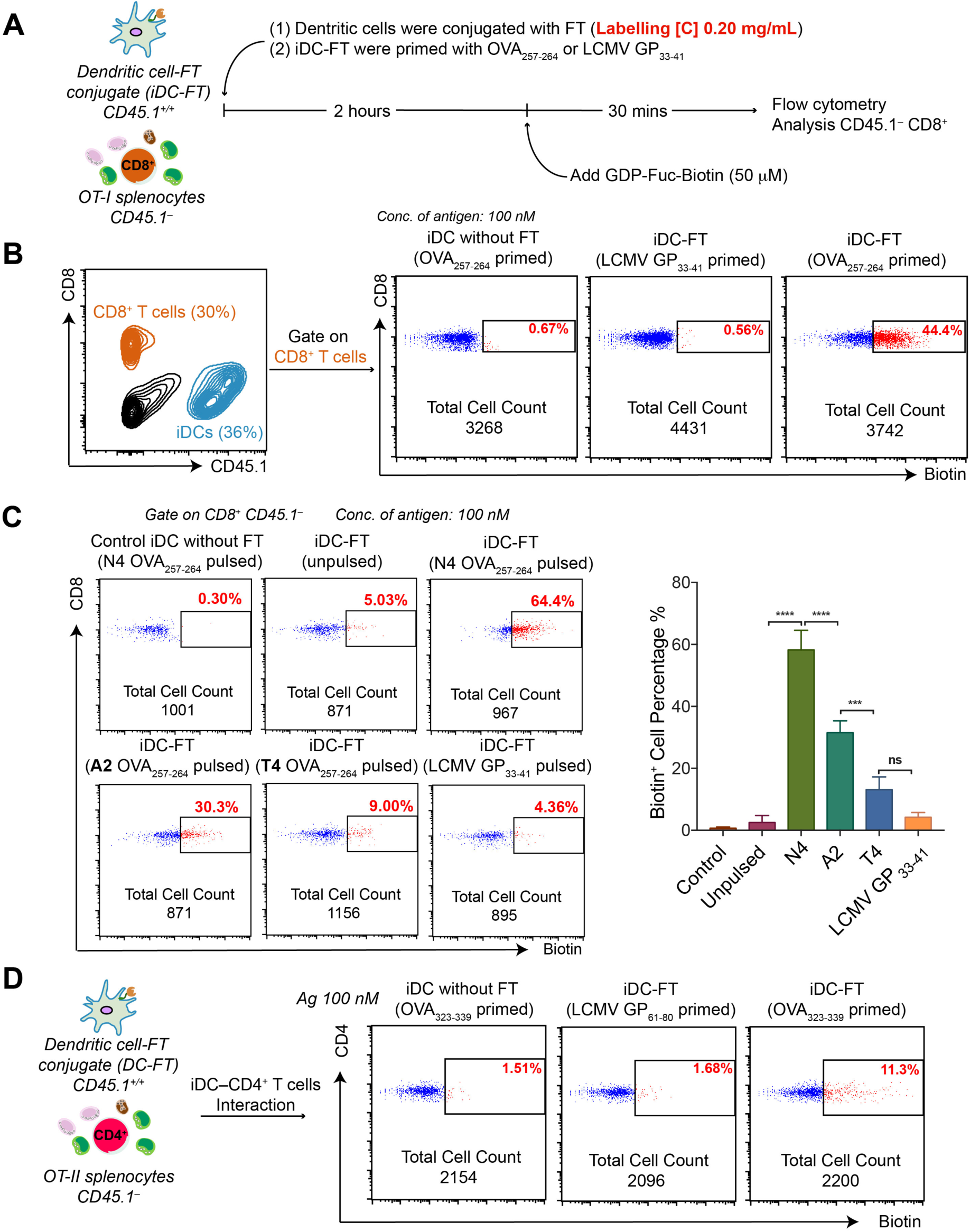
Characterization of the FucoID system in probing the interaction between iDCs and CD8^+^ or CD4^+^ T cells in splenocytes. Workflow for analyzing the interactions of antigen-primed iDCs with naïve OT-I T cells in OT-I splenocytes. **(B)** Flow cytometric analysis of antigen-specific Fuc-biotinylation of CD8^+^ T cells in OT-I splenocytes by iDC-FT loaded with OVA_257-264_ at the iDC/T cell ratio 1:1. **(C)** Flow cytometric analysis of OT-I CD8^+^ T cells Fuc-biotinylation by iDC-FT loaded with altered peptide ligands (APLs) derived from the original OT-I peptide SIINFEKL N4 (OVA_257-264_). mean ± SD (n=3); ns, P > 0.05; *P < 0.05; **P < 0.01; ***P < 0.001; ****P < 0.0001; one-way ANOVA followed by Tukey’s multiple comparisons test. **(D)** Flow cytometric analysis of CD4^+^ T cells specific Fuc-biotinylation by iDC-FT primed with OVA_323-339_ in OT-II splenocytes (iDC: T cell = 1:1). Representative figures from three independent replicates.

To explore if the labeling intensity achieved by FucoID reflects the strength of an interaction, we repeated the OT-I labeling experiment using iDCs primed with altered peptide ligands (APLs) derived from the original OT-I ligand SIINFEKL (N4). These APLs bind equally well to MHC-I H-2Kb as N4 but differ in their potency for interacting with TCR of OT-I CD8^+^ cells (binding strength: SII**N**FEKL(N4)>S**A**INFEKL(A2)>SII**T**FEKL(T4)) (Zehn et al., 2009). As shown by flow cytometry analysis (**Fig. 3C)**, the magnitude of intercellular labeling matched consistently with the TCR binding strength of the MHC-I bound APLs with the labeling induced by the N4-primed iDCs being the strongest (64.4%), which was followed by the labelling induced by iDC primed with A2 (30.3%), and the T4-primed iDC mediated labeling being the weakest (9.0%).

It was reported that surface molecules on APCs could be transferred to lymphocytes by trogocytosis. Although such trogocytosis is usually weak between primary immune cells, Baltimore et al. reported recently that trogocytosis of TCR proteins occurring from Jurkat-K562 interactions could be used to identify tumor neoantigen. In our framework settings, we confirmed that negligible amounts of Fuc-Bio or FT were transferred from iDC to the surface of T cells via trogocytosis (**Fig. S7**) (Li et al., 2019a).

Importantly, we found that FucoID could also be applied to probe iDC-CD4^+^ interactions despite significantly weaker binding between MHC-II bound peptides and CD4 (**Fig. 3D**). iDC-FT pulsed with OVA_323-339_ specifically biotinylated 11.3% of OT-II CD4^+^ T cells whose TCR was reactive with OVA_323-339_ under 2 hours of co-culturing. By contrast, only background labelling (1.68%) was observed in the group using LCMV GP_61-80_ pulsed DC-FT.

### Rapid detection and enrichment of tumor specific antigen (TSA) reactive TILs based on FucoID in a B16-OVA tumor model

With the validation of FucoID as a reliable technique for probing antigen-specific DC-T cell interactions *ex vivo*, we assessed the feasibility of using this strategy to detect and enrich TSA-reactive TILs from tumor digests. In our workflow, a harvested solid tumor is dissociated to prepare a single cell suspension and tumor lysates, in which the tumor lysates are used to prime autologous iDCs. Through this process, both non-mutant peptides and TSAs are loaded onto MHCs on the iDC surface. The primed iDCs are then subjected to FT conjugation and added to the single cell suspension, followed by the addition of GDP-Fuc-Biotin to initiate the interaction dependent fucosyl-biotinylation. Our hypothesis is that self-antigen-reactive T cells have already been eliminated in the thymus via “negative selection” (Klein et al., 2014; Xing and Hogquist, 2012); only T cells that interact with TSA-presenting iDCs would be labeled with Fuc-Bio **(Fig. 4A)**.

**Figure 4.**
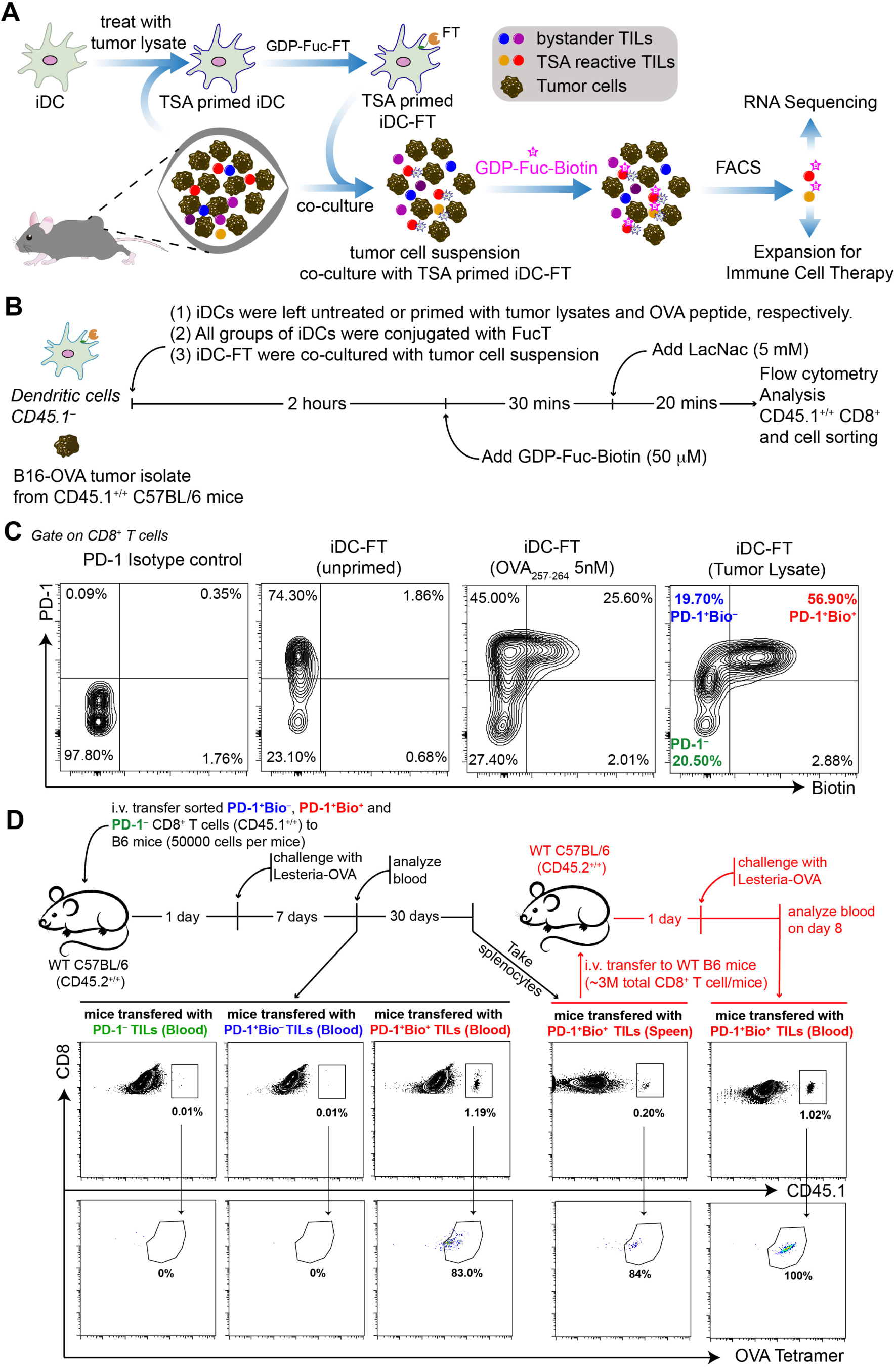
Identification of TSA-reactive T cells via FucoID from a B16-OVA melanoma model. (**A**) Illustration of the detection and enrichment of TSA-reactive TILs via FucoID. (**B**) Workflow of the FucoID system in identifying TSA-reactive TILs from a B16-OVA tumor. (**C**) Representative flow cytometric analysis of TSA dependent Fuc-biotinylation of CD8^+^ TILs in B16-OVA tumors. (**D**) Experimental design and flow cytometric analysis of the re-expansion and memory formation of the isolated PD-1^−^, PD-1^+^Bio^−^ and PD-1^+^Bio^+^ TILs after adoptive transfer into naïve mice upon OVA re-stimulation by LM-OVA; representative figures from three independent replicates.

The well-established B16-OVA melanoma was used as the first model system to test this hypothesis. In this model, the B16 melanoma cell line is engineered to express chicken ovalbumin as a TSA. Day 14 subcutaneous B16-OVA tumors were harvested individually from C57BL/6 mice (CD45.1^+/+^) and digested to prepare single cell suspensions for co-culturing with iDC-FT (CD45.1^−^) that were pre-treated with and without B16-OVA tumor lysates, respectively. GDP-Fuc-Bio was then added to initiate the interaction-dependent labeling. In this experiment, iDC-FT primed with OVA_257-264_ was used as the positive control (**Fig. 4B**). As revealed by flow cytometric analysis, whereas unprimed iDC-FT only labeled negligible numbers of CD8^+^ TILs (1.86%), iDC-FT primed by OVA_257-264_ or B16-OVA tumor lysates labeled 25.6% and 56.9% autologous CD8^+^ TILs respectively. Among all CD8^+^ TILs, ∼76% were PD-1^+^, within which approximately 74% were fucosyl-biotinylated by the iDC-FT primed with tumor lysates and according to our hypothesis this subset were bona fide TSA-reactive T cells. The remaining 26% of PD-1^+^ TILs were not labeled (**Fig. 4C**, last plot on the right), suggesting that this fraction may be bystander cells. Interestingly, only ∼3% PD-1^−^ T cells were labeled, indicating that almost all TSA-reactive T cell candidates had encountered with their cognate antigens.

To determine if the PD-1^+^Bio^+^ subset possesses better killing capabilities of B16-OVA cells than the PD-1^−^ and PD-1^+^Bio^−^ subsets, the labeled CD8^+^ TILs were FACS isolated. The isolated TILs were expanded using a reported rapid expiation protocol (Fernandez-Poma et al., 2017) and the tumor killing abilities were assessed on expansion day 7. As shown in **Figure S8A**, the expanded PD-1^+^Bio^+^ TILs showed significantly stronger killing of B16-OVA tumor cells than the other two subsets.

If bona fide TSA-reactive T cells were enriched in PD-1^+^Bio^+^ TILs, a sub-population of which should be OVA specific. To assess this, PD-1^+^Bio^−^ and PD-1^+^Bio^+^ TILs were cultured for 48 hours allowing the complete decay of the biotinylated molecules from the cell surface, at which point the cultured TILs were stained with H-2Kb/OVA_257-264_ MHC tetramer. Whereas the PD-1^+^Bio^+^ subset was found to contain 50% tetramer^+^ TILs, among which 6% stained tetramer ^high^, the PD-1^+^ Bio^−^ subset only contained 9.16% tetramer^+^ TILs and none of them were tetramer^high^ (**Fig. S8B**).

To determine if the isolated PD-1^+^Bio^+^, PD-1^+^Bio^−^ and PD-1^−^ subsets can undergo re-expansion and memory formation upon OVA stimulation, the isolated TILs (CD45.1^+/+^) were immediately transferred into antigen-free wild-type (WT) C57BL/6 (B6) hosts (CD45.2^+/+^) individually (**Fig. 4D**). One day later, these secondary recipients were immunized with *Listeria* expressing OVA (LM-OVA). Blood and/or spleens were isolated and analyzed on day 8 and day 38. Significant expansion of the transferred PD-1^+^Bio^+^ cells in blood were observed on day 8 (>1.1% of total blood CD8^+^) while no expansion of the transferred PD-1^+^Bio^−^ and PD-1^−^ subsets were detected (<0.05% of total blood CD8^+^) (**Fig. 4D**). Most of these expanded CD45.1^+^CD8^+^ T cells were H-2Kb/OVA_257-264_ tetramer positive (83%). On day 38, ∼0.2% of PD-1^+^Bio^+^ T cells remained detectable in the spleen and a majority of them were H-2Kb/OVA_257-264_ tetramer positive (84%). We then isolated the total CD8^+^ T cells from the spleens of these mice and transferred them into healthy B6 hosts (CD45.2^+/+^). The recipient mice were challenged with LM-OVA on the next day. On day 7 post infection, blood was collected and analyzed. We observed re-expansion of the transferred CD45.1^+/+^ CD8^+^ TILs in all recipient mice (∼1% of total blood CD8^+^) and all these expanded cells are OVA-tetramer positive (**Fig. 4D**). Together, these results provide solid support of FucoID as a highly effective approach to detect and enrich for TSA-reactive TILs from tumor cell suspensions for the subsequent adoptive transfer-based applications.

### TILs labeled via FucoID in multiple syngeneic murine tumor models are *bona fide* TSA-reactive TILs

After confirming FucoID as a highly effective approach to detect and enrich for TSA-reactive TILs from the B16-OVA melanoma model, we sought to explore its scope and limitation. Toward this end, tumor cell suspensions prepared from subcutaneous B16 melanoma, E0771 TNBC and MC38 colon tumors were subjected to the FucoID-based labeling (**Fig. 5A**). iDC-FT that were primed by tumor lysates fucosyl-biotinylated 34%, 17% and 25% autologous CD8^+^ TILs from B16 melanoma, E0771 TNBC and MC38 colon cancer, respectively **(Fig. 5B)**. By contrast, iDC-FT treated with the lysates obtained from the corresponding healthy tissues afforded only background labelling (<3%). Not surprisingly, iDC-FT primed with GP100_25-33_, a predominant human and B16 melanoma specific antigen, labeled 10% CD8^+^ TILs from the B16 tumor cell suspension. It is worth noting that in all three tumor models, almost all biotinylated TILs were PD-1^+^, suggesting that these TILs have encountered their cognate antigens. By contrast, 35%, 50% and 41% of TILs from each model were PD-1^+^Bio^−^, respectively, suggesting that they are irrelevant, bystander T cells, but displaying a different phenotype than PD-1^−^ bystander TILs.

**Figure 5.**
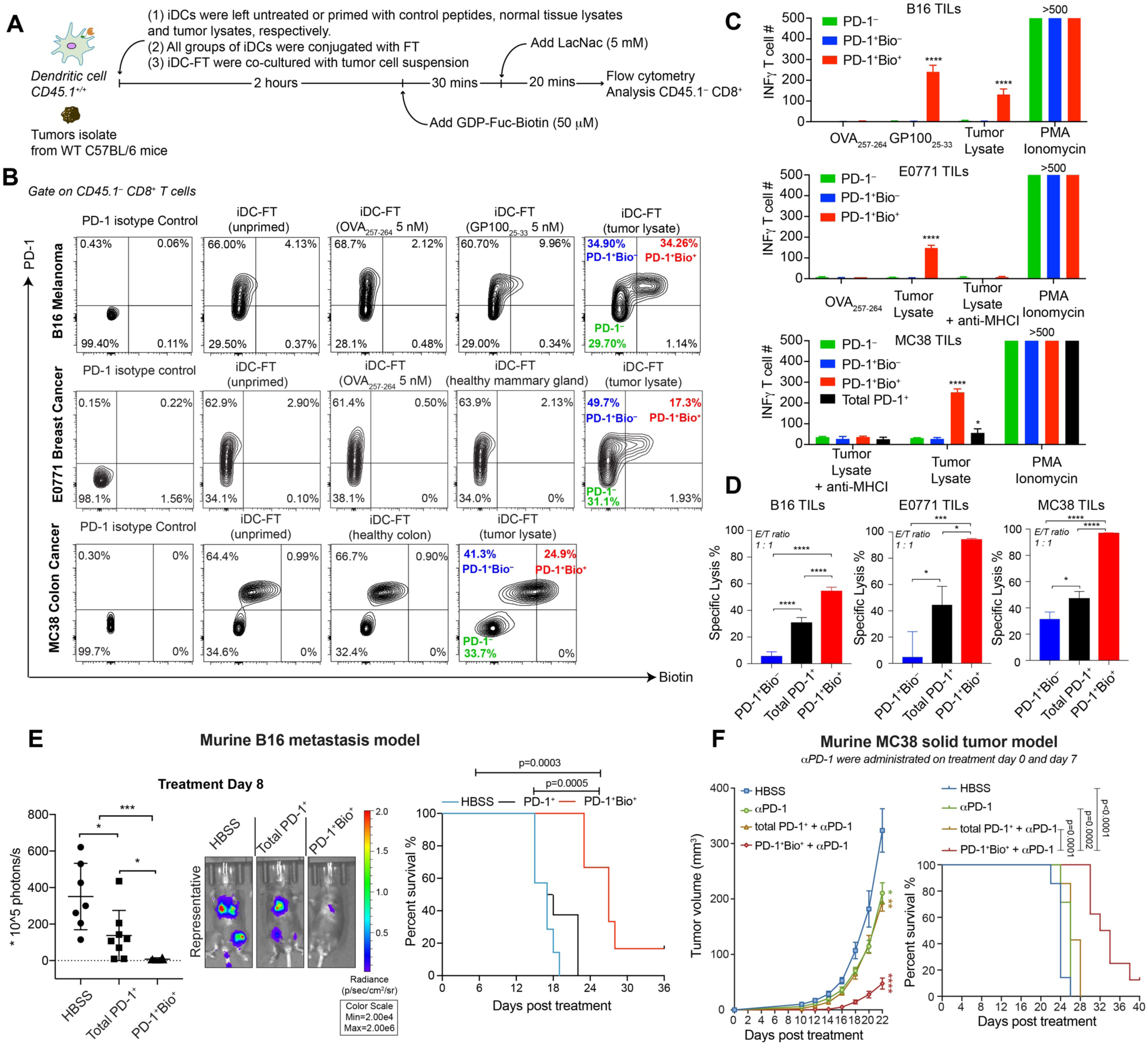
TSA-reactive CD8^+^ T cells identified and isolated from distinct murine tumor models via FucoID approach exhibit excellent tumor reactivity. (**A**) Workflow of the detection of TSA-reactive CD8^+^ T cells from syngeneic murine tumor models via FucoID. (**B**) Representative flow cytometric analysis of tumor antigen dependent Fuc-biotinylation of CD8^+^ TILs in B16 melanoma, E0771 TNBC and MC38 colon cancer models. (**C**) IFN-γ ELISPOT showing distinct reactivity of expanded PD-1^−^, PD-1^+^Bio^−^ and PD-1^+^Bio^+^ TILs upon tumor antigen re-stimulation. n=3; (**D**) Comparisons of specific lysis reactivity of expanded total PD-1^+^ TILs and PD-1^+^Bio^+^ TILs against the relevant cancer cells at effector-to-target ratio of 1:1; n=3; (**E**) In vivo tumor reactivity of PD-1^+^Bio^+^ and PD-1^+^ TILs in a murine B16 metastasis tumor model. C57BL/6 mice were intravenously injected with 0.5 × 10^6^ B16-luc cells. TILs treatments were performed as described in method on day 3 of tumor inoculation. On day 8 of TIL transfer, tumors were imaged by IVIS system. The sizes of the tumors and representative images are shown (mean ± SD); HBSS: 7 mice; total PD-1^+^: 8 mice; PD-1^+^Bio^+^: 6 mice; (**F**) In vivo tumor reactivity of PD-1^+^Bio^+^ and PD-1^+^ TILs in a murine MC38 s.c tumor model. MC38 cells were s.c. injected into the right flanks of male C57BL/6 mice (0.5 × 10^6^ cells per mice). Mice were irradiated (5 Gy) on day 2 of tumor inoculation. Then TILs treatments were performed as described in method on day 3 of tumor inoculation. Tumor volumes were measured every 2 days from day 10 of TILs treatment. The average size of tumors of each group until treatment day 22 are shown; HBSS: 7 mice; anti-PD-1: 7 mice; total PD-1^+^: 7 mice; PD-1^+^Bio^+^: 8 mice. In above figures, mean ± SD (error bars); ns, P > 0.05; *P < 0.05; **P < 0.01; ***P < 0.001; ****P < 0.0001; data were analyze by one-way ANOVA followed by Tukey’s multiple comparisons test or t-test. Survival data were analyzed by Log-rank (Mantel-Cox) test.

To characterize the function of these subsets, the isolated PD-1^−^, PD-1^+^Bio^−^ and PD-1^+^Bio^+^ TILs were cultured and expanded using the rapid expansion protocol in the presence of feeder cells, anti-mCD3 and rhIL-2 (**Fig 5B**). At day 9 upon expansion, we performed IFNγ ELISpot to assess the TSA reactivity of the expanded T cells. T cells were re-stimulated with DCs pulsed with tumor lysates. As the negative controls, T cells were re-stimulated by DCs pulsed with an irrelevant peptide. As expected, when re-stimulated with DCs primed with tumor lysates specific IFN-γ secretion was only detected in the expanded PD-1^+^Bio^+^ TILs, which was blocked by the addition of an anti-MHC-I antibody (**Fig. 5C** and **Fig. S10**).

The differences in TSA-reactivity of PD-1^+^Bio^+^ and PD-1^+^Bio^−^ CD8^+^ TILs would be resulted from the differences of their TCR clonotypic repertories. We characterized TCR clonotypic repertoires of these two TIL subsets isolated from E0771 and MC38 tumor models directly after their FACS isolation without further in vitro expansion. TCRβ deep sequencing was employed for quantifying the frequency of individual T-cell clonotype in each subset. A productive CDR3 sequence that does not contain stop codons or frame shifts represents a unique TCR clonotype, and the total number of unique sequences determines the clonal diversity in each subset, providing each subset has comparable total CDR3 reads. We found that TCRβs in the PD-1^+^Bio^+^ population were significantly more oligoclonal than their counterparts in the PD-1^+^Bio^−^ subset, suggesting the cells in the PD-1^+^Bio^+^ subset have undergone substantial TSA-driven clonal expansion (**Fig. S11)**. Furthermore, there was no overlap of the 10 most abundant TCRβ CDR3s found between these two subsets in either tumor models (**Fig. S11)**. The results from IFNγ ELISpot assay and TCRβ deep sequencing indicated that PD-1^+^Bio^+^ and PD-1^+^Bio^−^ TILs represent two functionally and clonotypically distinct T cell subsets that co-exist in the same tumors and share a certain degree of phenotypical similarities (e.g. PD-1^+^).

Because PD-1^+^ TILs consisted of not only TSA-reactive T cells but also bystander T cells, upon in vitro expansion, anti-tumor cytotoxicity of the entire PD-1^+^ TIL population should be considerably weaker than that of the PD-1^+^Bio^+^ TIL subset providing that TSA-reactive and bystander T cells share similar expanding rates. To assess this hypothesis, PD-1^+^Bio^+^, PD-1^+^Bio^−^ and total PD-1^+^ TILs isolated from the same tumors were subjected to the rapid expansion protocol. According to the recorded growth curve, PD-1^+^Bio^−^ and total PD-1^+^ TILs exhibited very similar expansion rate, which was significantly faster than that of PD-1^+^Bio^+^ TILs during the entire course of expansion (**Fig. S12**). These observations suggest that at the end of expansion bystander PD-1^+^Bio^−^ TILs become the dominant cell population within the expanded total PD-1^+^ TILs. On the expansion day 10, tumor killing capabilities of each subset were assessed. At different effector/target ratios PD-1^+^Bio^+^ TILs exhibited remarkably stronger tumor cell killing than the corresponding PD-1^+^Bio^−^ and total PD-1^+^ TILs (**Fig. 5D, Fig. S13**). These results suggest that due to the faster proliferation of bystander T cells within the entire PD-1^+^ TIL population, anti-tumor activities of total PD-1^+^ TILs become significantly weaker than those of the expanded TSA-reactive PD-1^+^Bio^+^ TILs at the end point of rapid expansion.

### The expanded PD-1^+^Bio^+^ TILs exhibit significantly higher anti-tumor activities than the entire PD-1^+^ TILs *in vivo*

To compare anti-tumor activities of the expanded PD-1^+^Bio^+^ TILs and the total PD-1^+^ subset in vivo, we first explored the use of the TILs isolated from the B16 melanoma model to control tumor growth in mice with established pulmonary micrometastases. Three days after the intravenous inoculation of B16 tumor cells stably transduced with firefly luciferase (B16-luc) to induce pulmonary metastasis, tumor-imbedded mice were treated with the expanded total PD-1^+^ TILs and PD-1^+^Bio^+^ TILs (3 × 10^6^ per mice), respectively, while the control group was injected with buffer only. Tumor proliferation was monitored by longitudinal, noninvasive bioluminescence imaging. As shown in **Fig. 5E**, the total PD-1^+^ TILs showed moderate therapeutic potency for preventing tumor proliferation with 60% lower bioluminescence than the HBSS control group on treatment day 8. In comparison, PD-1^+^Bio^+^ TILs significantly inhibited tumor growth, showing 98% lower bioluminescence signal than the HBSS control group. Remarkably, the survival of the tumor bearing mice was significantly elongated upon treatment with the PD-1^+^Bio^+^ TILs. All mice received buffer only and the total PD-1^+^ TILs died on treatment day 22 and 26, respectively. By contrast, all PD-1^+^Bio^+^ TIL recipient mice were alive until treatment day 27 and by the end of the experiment, still 20% mice remained alive (treatment day 40).

To assess capabilities of the expanded PD-1^+^Bio^+^ TILs of suppressing solid tumor growth we sought to use the TILs isolated from the subcutaneous MC38 tumors. Expanded total PD-1^+^ TILs and PD-1^+^Bio^+^ TILs (5 × 10^6^ per mice), respectively, were intravenously injected into mice with established subcutaneous MC38 tumors, followed by anti-PD-1 administration. In the control groups, mice were treated with anti-PD-1 and HBSS, respectively. Although anti-PD-1 and anti-PD-1 + total PD-1^+^ TIL treatments only showed modest tumor control, PD-1^+^Bio^+^ TILs combined with anti-PD-1 significantly slowed down tumor growth with median tumor size being only ¼ of that treated with anti-PD-1^+^ total PD-1^+^ TILs (treatment day 22, **Fig. 5F**). Moreover, whereas no mice in any control group survived up to day 28, 50% mice treated with PD-1^+^Bio^+^ TILs + anti-PD-1 were still alive by day 34. And one mouse was found to be tumor free by day 40 (**Fig. 5F**). These results indicated that the expanded PD-1^+^Bio^+^ TILs possess markedly higher activities to control tumor growth in vivo than the expanded total PD-1^+^ TILs that contain a large fraction of bystander T cells.

### PD-1^+^Bio^+^ TILs are distinct to PD-1^+^Bio^−^ TILs and displays activation/dysfunction gene signature

To gain an understanding of the genetic programs that underlie the phenotypical and functional features of the TSA-reactive and two different groups of bystander TILs, we characterized transcriptional profiles of PD-1^+^Bio^+^, PD-1^+^Bio^−^ and PD-1^−^ CD8^+^ T cells isolated from MC38 subcutaneous tumors. Principle component analysis (PCA) revealed that the transcript profiles of these three subsets of TILs shared substantial divergence (**Fig. 6A**). As identified by volcano plot messenger RNA (mRNA) comparisons between PD-1^+^Bio^+^ and PD-1^+^Bio^−^ CD8^+^ TILs, 290 transcripts were significantly upregulated or downregulated (**Fig. 6B**). By contrast, a total of 3704 genes were differentially expressed between PD-1^−^ and PD-1^+^Bio^−^ TILs (**Fig. S15**).

**Figure 6.**
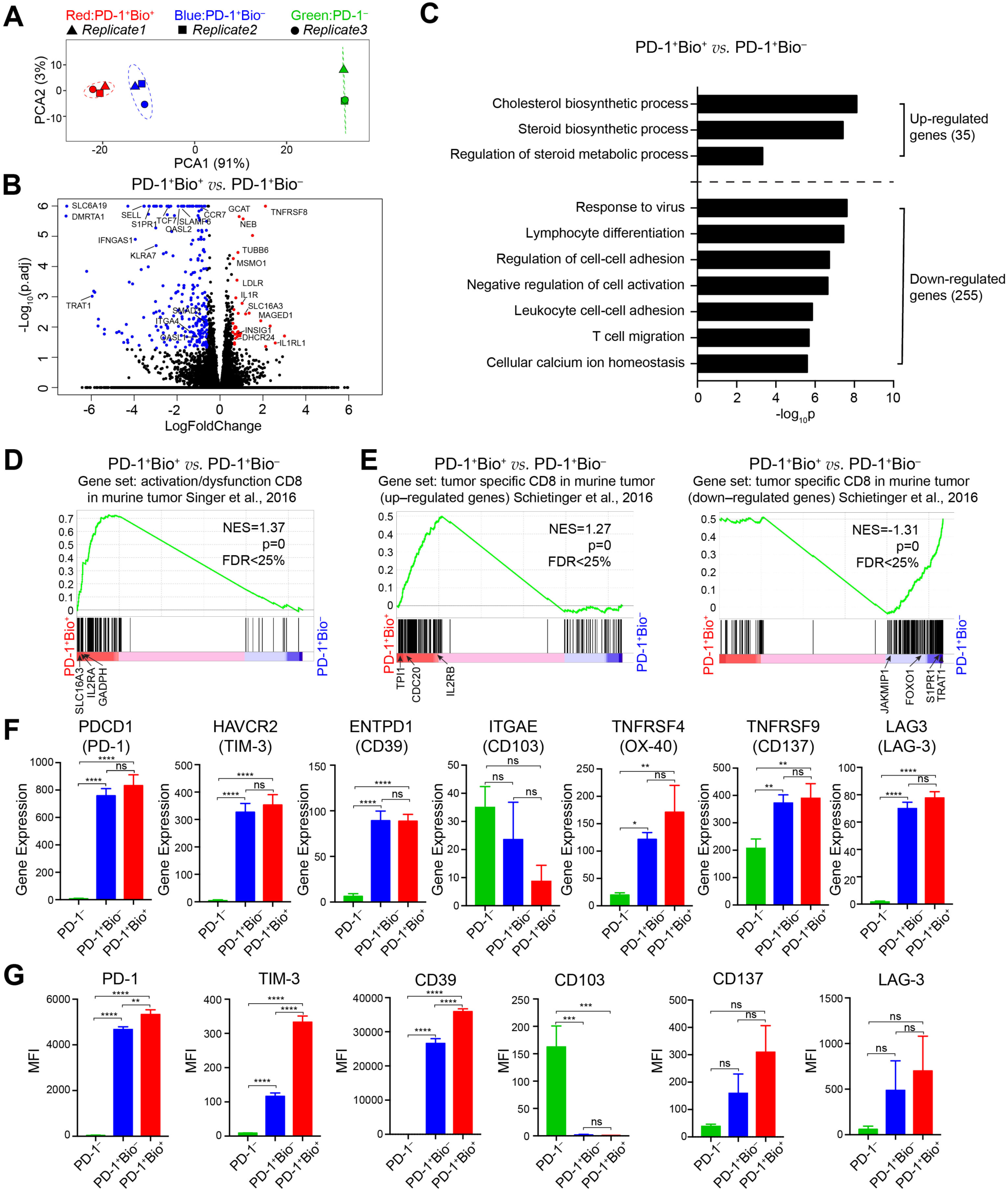
Distinct gene-expression signatures of PD-1^+^Bio^+^, PD-1^+^Bio^−^ and PD-1^−^ CD8^+^ TILs isolated from MC38 tumors. **(A)** PCA of the transcriptome of PD-1^+^Bio^+^ (red), PD-1^+^Bio^−^ (blue) and PD-1^−^ (green) CD8^+^ TILs isolated from murine MC38 tumors. Dots represent samples of the three different populations (grouped by colors) from a total of three biological replicates (grouped by shapes). PCA 1 and 2 represents the largest source of variation. (**B**) Volcano plot of up- (red) and down- (blue) regulated genes between PD-1^+^Bio^+^ and PD-1^+^Bio^−^ TILs. Significance was determined as Benjamini–Hochberg FDR (p.adjust) < 0.05 and |log_2_(fold change)| ≥ 0.6. (**C**) Biological processes (GO terms) enriched in the up– and down–regulated genes identified in **Figure 6B**. (**D**) Gene set enrichment analysis (GSEA) of activation/dysfunction CD8 gene module (Singer et al., 2016) in the transcriptome of PD-1^+^Bio^+^ *vs*. that of PD-1^+^Bio^−^ TILs. (**E**) GSEA of up- and down-regulated tumor specific CD8 gene signature (Schietinger et al., 2016) in the transcriptome of PD-1^+^Bio^+^ *vs*. that of PD-1^+^Bio^−^ TILs. n=3. (**F**) The comparison of representative gene expression of PD-1^+^Bio^+^, PD-1^+^Bio^−^ and PD-1^−^ TILs. (**G**) The expression of PD-1, CD137, TIM-3, CD39 and CD103 in PD-1^+^Bio^+^, PD-1^+^Bio^−^ and PD-1^−^ CD8 TILs from murine MC38 tumor according to flow cytometric analysis. Data obtained from at least two independent replicates. In above figures, mean ± SD (error bars); ns, P > 0.05; *P < 0.05; **P < 0.01; ***P < 0.001; ****P < 0.0001; data were analyze by one-way ANOVA followed by Tukey’s multiple comparisons test.

We then focused on analyzing the less pronounced transcriptional difference of PD-1^+^Bio^+^ and PD-1^+^Bio^−^ CD8^+^ TILs. An over-representation analysis was conducted to explore the enrichment of the 290 genes in biological processes annotated by the gene ontology database. Compared to PD-1^+^Bio^−^ TILs, several up-regulated genes of PD-1^+^Bio^+^ TILs were significantly enriched in steroid biosynthesis and related metabolic pathways (**Fig. 6C and S16**), such as MSMO1 and DHCR7. This is consistent with the previously reported discovery that the cholesterol metabolism of T cells is fully reprogrammed upon cell activation to support cell proliferation(Bensinger et al., 2008; Tuosto and Xu, 2018). An increase in the plasma membrane cholesterol level of CD8^+^ T cells augments T-cell receptor clustering, signaling and the more efficient formation of the immunological synapse, which are essential for the effector function of CD8^+^ T cells (Yang et al., 2016). By contrast, down-regulated genes are enriched in more diverse biological process networks, including those of lymphocyte differentiation, T cell migration and activation, viral response and calcium homeostasis (**Fig. 6C and S16**). These findings strongly suggest that PD-1^+^Bio^−^ CD8^+^ TILs are bystander T cells with virus reactivity and have an altered spectrum of core cellular processes compared to PD-1^+^Bio^+^ TILs (Scheper et al., 2019; Simoni et al., 2018).

To further characterize genetic differences of these three subsets of TILs, we performed gene set enrichment analysis (GSEA) using the gene signatures and gene modules established in chronic virus infection induced T cell exhaustion and tumor associated T cell activation/dysfunction models. We initially compared these three subsets TILs for the enrichment of a previously reported naïve/memory-like T cell gene module and found this module is enriched in PD-1^−^ TILs (**Fig. S17A and Table S1**) (Singer et al., 2016). Next, we assessed the three subsets TILs for the enrichment of the exhaustion signature derived from exhausted T cells isolated from chronic LCMV infection TILs (Wherry et al., 2007), finding a similar enrichment of this signature in both PD-1^+^Bio^+^ and PD-1^+^Bio^−^ CD8^+^ subsets compared to PD-1^−^ TILs (**Fig. S16B and Table S1**). We then compared the TSA-reactive PD-1^+^Bio^+^ TILs with the bystander PD-1^+^Bio^−^ and PD-1^−^ TILs for the enrichment of the gene modules shared by T cells infiltrating human or murine tumors. The T cell activation/dysfunction gene module established for B16F10 melanoma(Singer et al., 2016) was significantly enriched in PD-1^+^Bio^+^ *vs*. PD-1^+^Bio^−^ TILs; notable genes in this module include genes encoding the cytokine IL2 receptor (IL2RA), the T cell activation related glycolysis enzyme GAPDH (GAPDH) and the plasma membrane transporter for monocarboxylates such as lactate and pyruvate (SLC16A3) (**Fig. 6D, S17C and Table S1**). Consistent with this finding, an enrichment of the upregulated cell cycle gene signature that was validated for human melanoma TILs was found in PD-1^+^Bio^+^ TILs in comparison to both PD-1^+^Bio^−^ and PD-1^−^ bystander TILs(Li et al., 2019b) (**Fig. S17D and Table S1**). This finding, combined with the observed clonal expansion of PD-1^+^Bio^+^ TILs, provided strong evidence for ongoing proliferation within this dysfunctional but TSA-reactive T cell subset. This feature has also been observed previously for a subpopulation of CD8^+^ T cells infiltrating human and murine tumors that are believed to possess tumor reactivity. By comparing the transcript profiles of monoclonal CD8^+^ T cells specific for Tag epitope I (Tag-I; SAINNYAQKL) infiltrating early and late stage murine tumors to that of D30 exhausted T cells isolated from chronic LCMV infection, Greenberg and coworkers discovered the unique gene signatures of dysfunctional, tumor-specific T cells that are not shared by dysfunctional T cells triggered by chronic viral infection(Schietinger et al., 2016). We compared our polyclonal TSA-reactive PD-1^+^Bio^+^ TILs with the bystander PD-1^+^Bio^−^ and PD-1^−^ TILs for the enrichment of this tumor-specific T cell gene set and found that it was significantly enriched in the PD-1^+^Bio^+^ subset (**Fig. 6E, Fig. S17E and Table S1**).

Finally, we analyzed the transcript levels of the genes that were previously reported as tumor reactive TILs selection markers in PD-1^+^Bio^+^ and PD-1^+^Bio^−^ TIL subsets isolated from the MC38 colon cancer model. To our surprise, in both TIL subsets similar transcript expression levels were detected for all of these selection markers, including PDCD1 (PD-1), HAVCR2 (TIM-3), LAG3 (LAG-3), ENTPD-1 (CD39), ITAGE (CD103), TNFRSF4 (OX-40) and TNFRSF9 (CD137) (**Fig. 6F**). We further analyzed the expression of several of these selection markers on the cell surface by flow cytometry (**Fig. 6G and S18**). Although the PD-1^+^Bio^+^ TIL subset was found to express higher levels of TIM-3 and CD137 than the PD-1^+^Bio^−^ TILs, varying levels of LAG-3, CD39, and CD103 expressions were found in both subsets(Duhen et al., 2018; Gros et al., 2014a; Yossef et al., 2018). Taken together, we conclude that FucoID may be more generally applicable than these previously reported functional markers-based selection approaches to identify TSA-reactive TILs.

## Discussion

TILs within individual tumors consist of heterogeneous populations including not only the T cells specific for tumor antigens, but also those recognizing a wide range of epitopes unrelated to cancer (e.g. antigens from Epstein–Barr virus, human cytomegalovirus or influenza virus) (Scheper et al., 2019; Simoni et al., 2018). These bystander CD8^+^ TILs have diverse phenotypes that overlap with those of the tumor-specific T cells, but are not tumor reactive (Duhen et al., 2018; Yossef et al., 2018). Although several selection markers (e.g. PD-1, CD39, CD103) have been utilized to exclude bystander CD8^+^ TILs, they are empirical and may generate false positive selection. Moreover, TSA reactive TILs in less abundant or rare populations could be missed using such indirect selection methods because these TILs may not share the same exhaustion status or phenotypes with the most abundant TSA-reactive TILs. By contrast, the FucoID strategy developed here generates a selection maker, i.e. biotin, based on the direct TCR-pMHC interaction, thus providing an unbiased approach for TSA-reactive TIL identification. Through FucoID, although we would not be able to elucidate the identity of TSAs, T cell candidates that are TSA-reactive would be enriched directly for expansion and for rapid isolation of the corresponding TCRs to construct TCR engineered T cells for functional evaluation. As a consequence, research time and cost are dramatically reduced compared to the aforementioned reverse immunology based pMHC tetramer approach(Arnaud et al., 2020). Significantly, once a TSA-reactive TCR is confirmed, it is possible to use other recently developed methods to identify the corresponding antigen (Gee et al., 2018; Kula et al., 2019; Li et al., 2019a).

TSA-reactive CD8^+^ T cells (i.e. PD-1^+^Bio^+^ T cells) isolated by FucoID from murine tumor models in this study exhibited a dysfunctional phenotype, but still possessed significant proliferative and tumor killing capacities. A subpopulation of these TSA-reactive T cells (∼5%) harbored progenitor exhausted T cell characteristics (TCF1^+^TIM-3^*–*^) (**Fig. S18B**), which is in line with tetramer-sorted tumor-specific T cells from previous studies (Miller et al., 2019). It has been demonstrated that it is this subset of CD8^+^ T cells that provides the proliferative burst and effector function following anti-PD-1/PD-L1 therapy (Held et al., 2019; Im et al., 2016; Miller et al., 2019; Siddiqui et al., 2019). Therefore, future efforts should be devoted to approaches for enlarging this subset during rapid expansion to boost the therapeutic potential of TIL-based adoptive cell transfer.

We demonstrated here that FT modified mouse DCs could induce antigen specific fucosyl-biotinylation of not only CD8^+^ but also CD4^+^ T cells, and the labeling strength was correlated to the binding affinities of pMHC to TCR. Thus, FucoID opens a new door to study primary antigen specific CD4^+^ T cells not only in tumors but also in autoimmune diseases. CD4^+^ T cells are challenging targets to study using conventional approaches partially due to the diversity and length variation (11 to 30 amino acids) of MHC-II binding epitopes and their weak interactions with MHC-II (Editorial, 2017; Racle et al., 2019). Likewise, one can envisage applying FucoID to separate T cells possessing high affinity TCRs from those with weaker ones for studying their functions in tumor and related infection models.

As the first glycosyltransferase-mediated tagging approach for probing cell-cell interactions, FucoID does not rely on genetic manipulations such that it is readily applicable to probe primary cell interactions. Importantly, installing FT onto human DCs is as easy and straightforward as what has been shown here for human DC functionalization (**Fig. S2A**). Therefore, FucoID has a high potential to be translated to a clinical setting for the detection and isolation of TSA-reactive TILs from human patients. Through popularizing FucoID, we expect the pace for the discovery of TSA-reactive TILs and their TCRs would be significantly accelerated, which in turn would pave the way for lowering the cost and accessibility of personalized cancer treatment (Arnaud et al., 2020; Yamamoto et al., 2019).

## Supporting information

Table S1. GSEA ranking gene lists

## Author Contributions

Z.L., J.L. and M.C. contributed equally to this work. P.W., J.L. and Z.L. designed the experimental strategies and wrote the manuscript. Z.L., M.C., J.L. Y.S. and M.W. performed experiments. P.W., Z.L. and J.L. prepared the figures, and all authors analyzed the data and edited the manuscript.

## Notes

The authors declare the following competing financial interest(s): Z.L., J.L. J.R.T. and P.W. are listed as inventors on a patent application filed on June 3, 2019 (U.S. Patent Application No.: 62/856,551). All experiments were performed at Scripps Research, La Jolla.

## Acknowledgement

This work was supported by the NIH (AI143884 to P.W. and J.R.T; GM093282 to P.W.), we thank for Dr. Rechard Klemke (UCSD, USA) for E0771 cancer cells, Prof. James C. Paulson (Scripps Research, USA) for OT-II mice, Dr. Brunie H. Felding for IVIS imaging system and NIH tetramer facility for H-2Kb/OVA_257-264_ MHC tetramer.

## STAR★METHODS

### KEY RESOURCES TABLE

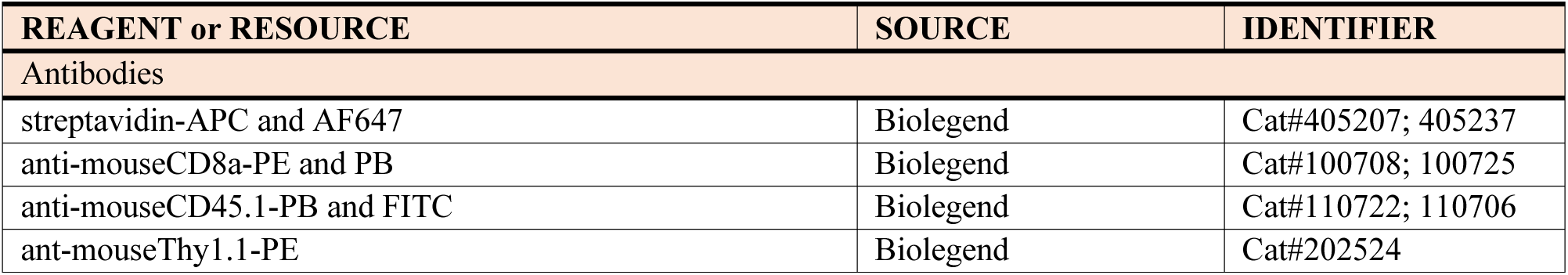
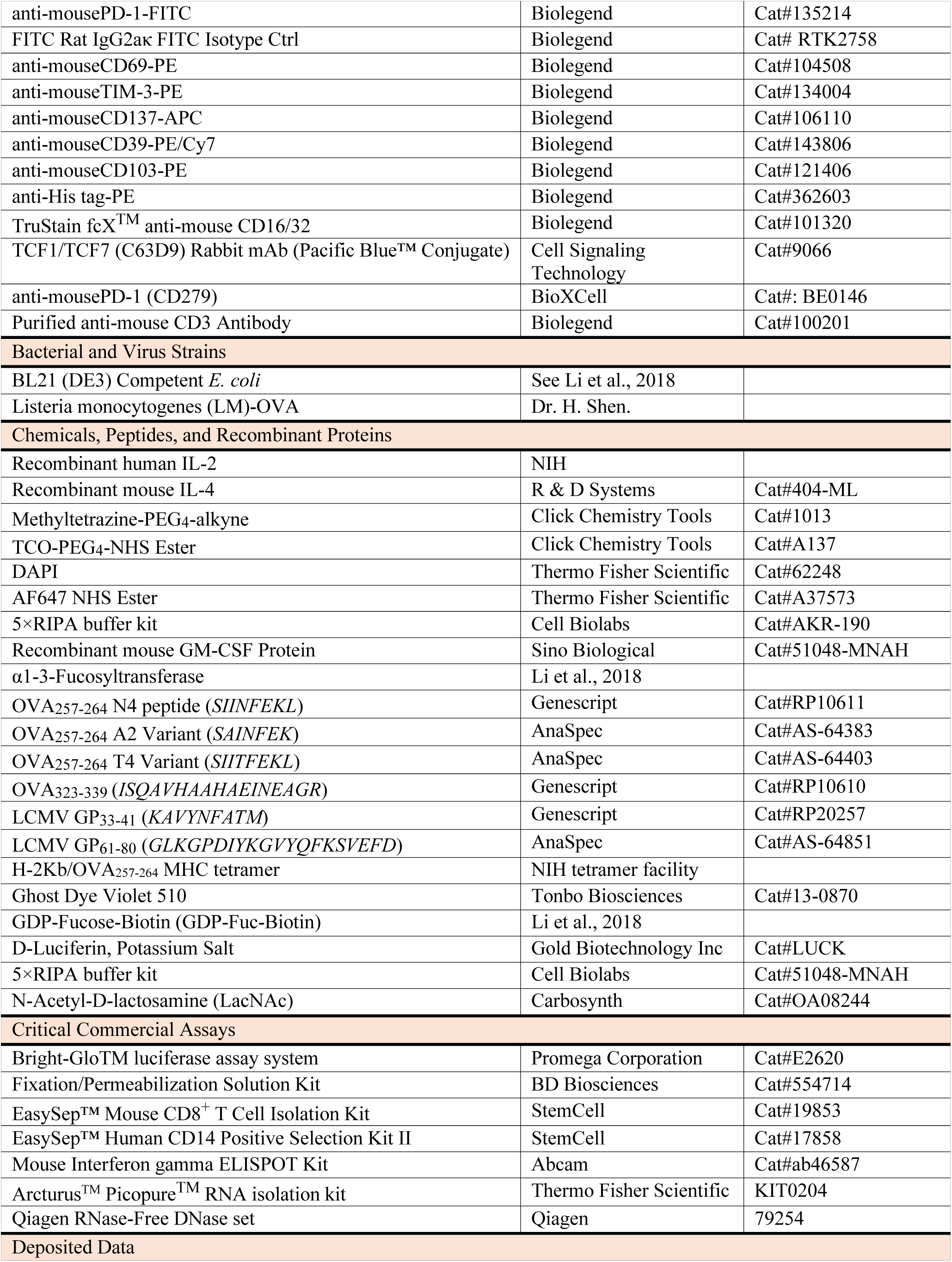
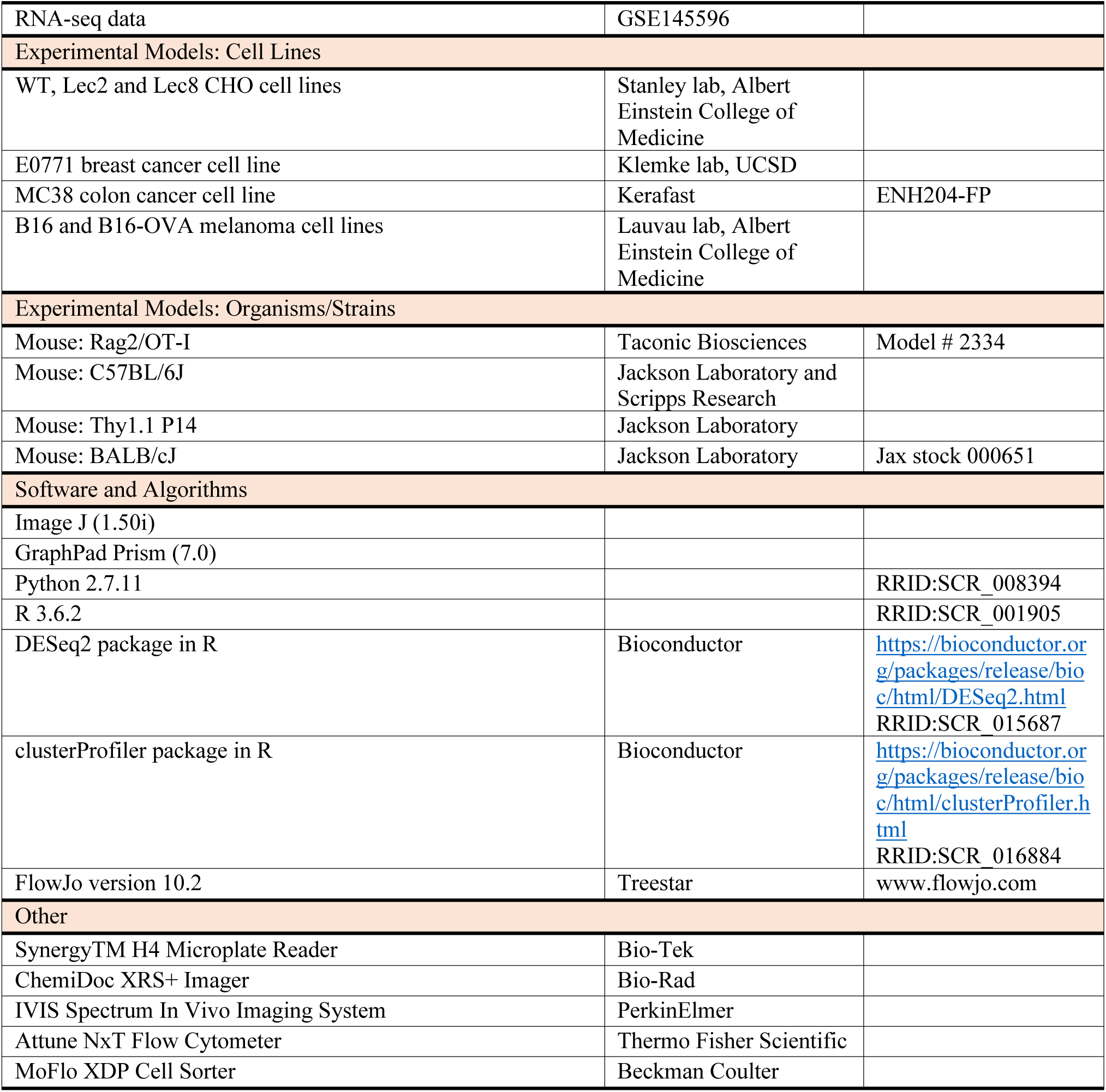

## CONTACT FOR REAGENT AND RESOURCE SHARING

**Further information and requests for resources and reagents should be directed to and will be fulfilled by the Lead Contact, Peng Wu**

