## supplementary information for "Detecting Tumor Specific Antigen-Reactive T cells from Tumor Infiltrating Lymphocytes via Interaction Dependent Fucosyl-biotinylation"

**EXPERIMENTAL MODEL AND SUBJECT DETAILS**

*Primary OT-I T cells preparation*

*Immature bone marrow dendritic cells (iDCs) and antigen priming*

*Human peripheral blood lymphocytes*

*Mice*

**Methods**

*1. One-pot protocol for producing GDP-Fucose-1,3-fucosyltransferase (GDP-Fuc-FT)*

*2. General protocol for enzymatic transfer of GDP-Fuc-FT to cell surface*

*3. Fluorescent imaging of the FucoID enabled proximity labelling on CHO cells*

*4. Detection of iDC–OT-I CD8$^+$ T cell interaction by FT functionalized dendritic cells (DC-FT)*

*5. Detection of selective interaction of dendritic cells in the mixture of OT-I CD8$^+$ T cells and P14 CD8$^+$ T cells*

*6. Identification and enrichment of TSA reactive TILs via FucoID*

*7. In vivo re-stimulation of different subsets of TILs from B16-OVA tumor*

*8. Rapid expansion of TILs*

*9. IFN$\gamma$ EliSpot assay for TILs reactivity analysis*

*10. In vitro tumor killing assays of TILs*

*11. TCR$\beta$ sequencing, mRNA sequencing and analysis*

*12. Generation of gene signatures from the literature*

*13. Statistical analysis*

**Supplementary Figures**

*Figure S1. Synthesis of GDP-Fuc-Biotin and GDP-Fuc-Tz*

*Figure S2. Conjugation of the recombinant FT onto CHO cell surface.*

*Figure S3. Conjugation of FT onto multiple types of cells.*

*Figure S4. Comparison of the antigen presentation abilities of iDC and iDC-FT*

*Figure S5. Fluorescent imaging of FucoID mediated intercellular labeling of CHO cells.*

*Figure S6. Optimization of FucoID condition in the iDC-OT-I CD8$^+$ T co-culturing system.*

*Figure S7. Trogocytosis is not observed during antigen-dependent intercellular labeling via FucoID*

*Figure S8. Tumor reactivity and TCR specificity of PD-1$^+$Bio$^-$ and PD-1$^+$Bio$^+$ TILs isolated from murine B16-OVA tumor via FucoID.*

*Figure S9. Representative flow cytometry figures showing the gating strategy of E0771 TILs after FucoID labelling, related to Figure 5B.*

*Figure S10. Representative images of IFN$\gamma$ EliSpots of TILs, related to Figure 5C.*

*Figure S11. Representative TCR$\beta$ RNA sequencing results of PD-1$^+$Bio$^-$ and PD-1$^+$Bio$^+$ TILs from E0771 tumor and MC38 tumor.*

*Figure S12. Cell expansion curves of total PD-1$^+$, PD-1$^+$Bio$^-$ and PD-1$^+$Bio$^+$ TILs under the rapid expansion protocol.*

*Figure S13. Comparisons of the cancer cell killing activity of expanded total PD-1$^+$ TILs and PD-1$^+$Bio$^+$ TILs at different effector-to-target cell ratios.*

*Figure S14. Individual tumor growth of the MC38 model with adoptive TILs transfer treatment, related to Figure 5F.*

*Figure S15. Volcano plot of up- (red) and down- (blue) regulated genes of PD-1$^+$Bio$^-$ vs. PD-1$^-$ TILs.*

*Figure S16. Enriched biological process by up- and down-regulated genes in PD-1$^+$Bio$^+$ vs. PD-1$^+$Bio$^-$ TILs, related to Figure 6C.*

*Figure S17. Gene set enrichment analysis (GSEA) of the subsets of MC38 TILs using reported gene signatures, related to Figure 6D and E.*

*Figure S18. Flow cytometric analysis of three subsets of CD8$^+$ TILs from murine MC38 tumor, related to Figure 6G.*

### EXPERIMENTAL MODEL AND SUBJECT DETAILS

#### Cell Lines

Cell lines were all purchased from ATCC unless otherwise specified. CHO cell lines (WT, Lec2 and Lec8, from Prof. Pamela Stanley at Albert Einstein College of Medicine) were grown as monolayer in alpha-Minimum Essential medium (α-MEM) (GIBCO) supplemented with 10% fetal bovine serum (FBS) (Omega Scientific, Inc). Cancer cell lines including E0771 (from Dr. Klemke lab, UCSD), MC38 (Kerafast), mouse B16 and B16-OVA (from Prof. Gregoire Lauvau lab) are all grown in DMEM (Dulbecco's modified Eagle's medium, GlutaMAX, GIBCO) supplemented with 10% FBS. All cells cultures were incubated at 37 °C under 5% $CO_2$.

#### Primary OT-I T cells preparation

Naïve OT-I CD8$^+$ T cells or splenocytes were isolated from the spleen of OT-I mice with or without the mouse CD8$^+$ isolation kit. T cells or splenocytes were cultured in RPMI 1640 (GlutaMAX) with 10% heat-inactivated FBS, 1 mM sodium pyruvate, 50 μM β-ME, 10mM HEPES and 1×MEM NEAA, 100 U/ml penicillin and 100 mg/ml streptomycin (referred as complete T cell culture media later). Cytokines in T cell culture media were added as indicated. To generate effector CD8$^+$ T cells, OT-I splenocytes were cultured in complete T cell media containing 1 nM SIINFEKL ($OVA_{257-264}$) peptide for 2 days, followed by supplying 100 IU/mL rhIL-2. After culturing for additional 4 days, effector $CD8^+$ T cells were ready for use. All cells cultures were incubated at 37 °C under 5% $CO_2$.

#### Immature bone marrow dendritic cells (iDCs) and antigen priming

Bone marrow was taken from $CD45.1^{+/+}$ or WT ($CD45.2^{+/+}$) C57BL/6J mice. Following erythrocyte lysis, the bone marrow cells were resuspended in complete T cell medium with GM-CSF (20 ng/mL) and rmIL-4 (5 ng/mL). The culture medium was changed every 3 days with fresh complete T cell medium with GM-CSF (20 ng/mL) and rmIL-4 (5 ng/mL). After 7 days culturing, non-adherent cells and loosely adherent cells were harvested and gently washed by PBS for subsequent labelling experiments.

For the experiments of studying iDC-OT-I $CD8^+$ T cell interactions, iDCs or FT functionalized iDCs were cultured with or without indicated antigen peptide in complete T cell medium at 37 °C for 30 min. Then non-adherent cells in the culture supernatant and loosely adherent cells were harvested and gently washed with PBS for subsequent experiments.

For the experiments of labelling TSA-reactive TILs and IFNγ EliSpot assay, iDCs were cultured with or without indicated antigen peptide or tumor lysates (tumor cell/DC ratio = 10:1) in complete T cell medium containing GM-CSF (20 ng/mL) for 16 hours. Then non-adherent cells in the culture supernatant and loosely adherent cells were harvested and gently washed with PBS for subsequent experiments.

#### Human peripheral blood lymphocytes

Human blood samples were collected from healthy donors under the TSRI Normal Blood Donor Services program (#IRB 15-6710). Peripheral blood mononuclear cells (PBMCs) were obtained by Ficoll (Ficoll-Paque Plus, GE) density centrifugation.

*Primary human T cells preparation*

4 Million per mL PBMCs were cultured in T cell culture media with 15 ng/mL rhIL-2 and activated with human CD3/CD28 T cell activator for two days. After that, activated human T cells were kept under $4\times10^6$ cells/mL in T cell culture media (fresh media with cytokine were added every two days). Phenotypes were characterized after two weeks expansion (>95% are $CD3^+$ human T cells). The cells were then used for FT labelling.

*Primary human DC preparation*

$CD14^+$ monocytes were isolated from PBMCs using EasySep™ Human CD14 Positive Selection Kit II. Then, the CD14+ monocytes were treated with rhIL-4 (10 ng/mL) and recombinant human GM-CSF (20 ng/mL) by pipetting the cytokines directly into the complete T cell medium, incubate at 37 °C and 5% $CO_2$. Immature human DCs were harvest for FT labelling on day 7.

#### Mice

All mice were bred or housed under specific pathogen free (SPF) conditions. All animal experiments were approved by TSRI Animal Care and Use Committee. OT-I mice were purchased from Taconic Biosciences. $CD45.1^{+/+}$ and $Thy1.1^{+/+}$ mice from C57BL/6J genetic background were purchased from the Jackson Laboratory. The strain of $OT\text{-}I^{+/-}CD45.1^{+/-}$ and $OT\text{-}I^{+/-}Thy1.1^{+/-}$ was generated by cross breeding. Both male and female mice of 8–15 weeks of age were used for most experiments.

**Tumor Experiments**

*Tumor inoculation for TILs isolation*

1 x $10^6$ B16, B16-OVA or MC38 tumor cells were implanted subcutaneously into the flanks of male C57BL/6 mice (CD45.1$^-$ if no specific indication). 1 x $10^6$ E0771 tumor cells were implanted subcutaneously into the breast of female C57BL/6 mice. Fifteen days later, tumors were excised for use.

*Tumor inoculation for assessment of the tumor reactivity of TILs in murine tumor models*

*B16 tumor model*

Autologous CD8$^+$ TILs were labelled and sorted from B16 tumor (excised from male C57BL/6 mice) based on FucoID as shown in Figure 5. Then, TILs were subjected to the rapid expansion condition and ready for use on expansion day 10. Meanwhile, B16-luc (stably transduced with firefly luciferase) tumor cells (0.5 x $10^6$ per mice) were implanted subcutaneously to male C57BL/6 mice by tail vein injection. Three days later, mice were injected with 200 μL D-luciferin (15mg/mL) through intraperitoneally injection and tumor size were imaged according to the bioluminescence signal. The mice were then divided into 3 groups to make sure the initial tumor sizes were even in each group. Then, each group were treated with HBSS, total PD$^+$ TILs or PD-1$^+$Bio$^+$ TILs by tail vein injection (3 x $10^6$ per mice), respectively. After treatment, mice were i.p. injected with 50000 IU/mice rhIL-2 every 12 hours for 4 days. On day 8 of TILs treatment, mice were injected with 200 μL D-luciferin (15mg/mL) through i.p. injection. 10 minutes later, the bioluminescence signal in mice were analyzed by PerkinElmer IVIS system. The total photons indicating the tumor mice were quantified by IVIS software. Survival of mice was tracked from day 0 of TILs treatment to day 36 of TILs treatment.

*MC38 tumor model*

Autologous CD8$^+$ TILs were labelled and sorted from MC38 tumor (excised from male mice) based on FucoID as shown in Figure 5. Then TILs were subjected to the rapid expansion condition and ready for use on expansion day 10. Meanwhile, MC38 tumor cells (0.5 x $10^6$ per mice) were s.c. injected into the right flanks of male C57BL/6 mice. Two days later, all mice were irradiated (5 Gy). On the next day, mice were randomly divided into 4 groups and were treated with HBSS, anti-PD-1 (i.p., 100 μg per mice), PD-1$^+$ TILs (i.v., 5 x $10^6$ per mic) + anti-PD-1 (i.p., 100 μg per mice) or PD-1$^+$Bio$^+$ TILs (i.v., 5 x $10^6$ per mic) + anti-PD-1 (i.p., 100 μg per mice). After treatment, mice were i.p. injected with 50000 IU/mice rhIL-2 every 12 hours for 4 days. Another dose of anti-PD-1 (i.p., 100 μg per mice) was given to all mice except the HBSS control group on day 7 post treatment. Tumor major axis D and minor axis d were measured every 2 days from day 10 post treatment. Tumor volume ($mm^3$) = 1/2 x D x $d^2$. Survival of mice was tracked from day 0 of TILs treatment to day 40 of TILs treatment. A mouse was considered as 'death' when its tumor volume exceeded 400 $mm^3$.

#### Methods

##### 1. One-pot protocol for producing GDP-Fucose-1,3-fucosyltransferase (GDP-Fuc-FT)

Reactions were typically carried out in a 1.5 mL Eppendorf tube. TCO group was first introduced onto α(1,3)-fucosyltransferase (FT, see ref for expression and purification protocol) according to the standard labeling protocol of TCO-$PEG_4$-NHS ester (https://clickchemistrytools.com/product/tco-peg4-nhs-ester/) and previous reports (ref) about its application on IgG labeling. Briefly, we prepared fresh 50 mM stock of TCO-$PEG_4$-NHS reagent in DMSO and add it to the FT protein solution (5 mg/mL, 200 μL) at a final concentration of 2 mM. The reactions were incubated at room temperature for 30 minutes and quenched by adding

Tris buffer (pH 8.0, final concentration of 50mM). The quenched reaction mixtures were incubated at room temperature for 5 minutes and then desalted into PBS using G25 desalting column (PD-10, GE). The concentration of desalted TCO-FT was ~4 mg/mL (molecular weight distribution was confirmed in Fig.S2). After that, ~10 mM one-pot products of GDP-Fuc-Tz (ref) were added to TCO-FT with a final concentration at 0.15 mM (5 equiv. of TCO-FT). After 2 hours of incubation at room temperature, these one-pot GDP-Fuc-FT were ready to use and could be kept at -20 °C for up to 2 months.

**2. General protocol for enzymatic transfer of GDP-Fuc-FT to cell surface**

1 x $10^6$ Live cells were resuspended in 100 µL HBSS buffer containing 20 mM $MgSO_4$, 3 mM HEPES and 0.5% FBS. The cells were then treated with 0.2 mg/mL GDP-Fuc-FT (other concentrations were used in the titration experiment). With the reaction for 20 minutes on ice, cells were washed with PBS twice and ready for further application or analysis. For flow cytometry analysis, FT labeled cells were stained with DAPI and fluorescent secondary antibody (anti-His tag-PE) against labeled FT (His tag recombinant protein) on ice for 30 minutes. For the SDS-PAGE fluorescent gel imaging analysis, GDP-Fuc-FT was prelabelled with fluorescent dye AF647 (GDP-Fuc-FT-AF647) using AF647 NHS Ester and used instead of GDP-Fuc-FT. After cells were labelled with GDP-Fuc-FT-AF647 (0.2 mg/mL) on ice for 30 mins, labeled cells or untreated cells were collected and washed three times. After that, counted cells were lysed in SDS loading buffer and subjected to SDS-PAGE analysis. Quantitative GDP-Fuc-FT-AF647 protein were used as standards for quantification. For example, $1\times10^5$ CHO cells labeled with FT-AF647 were lysed in 20 µL SDS loading buffer and loaded to one lane of the SDS-PAGE gel. Meanwhile, 1 ng, 3 ng and 9 ng GDP-Fuc-FT-AF647 protein were loaded on the same page gel as the references. The resolved fluorescent gel was analyzed by ChemiDoc XRS+ (Bio-Rad). According to the semi-quantitative experiment, each CHO cell has $9 \times 10^5$ FT molecules. Similarly, each iDCs has 5 x $10^5$ FT molecules. (see Li. et al. 2018 for quantitative and calculation details)

**3. Fluorescent imaging of the FucoID enabled proximity labelling on CHO cells**

CHO cells pre-stained with CFSE were treated with 0.2 mg/mL GDP-Fuc-FT to generate FT functionalized CHO cells (CHO-FT). CHO-FT were mixed with untreated CHO cells at the ratio of 1:5 and the cell mixtures were allowed to rest in a glass chamber (Thermal Fisher Scientific). Cells were incubated at 37 °C for 2 hours in αMEM with 20 mM $MgSO_4$, then GDP-Fuc-biotin (ref, 50 µM) were added. After 20 min, medium was removed, and the glass chamber was gently washed twice with PBS and then cells attached on the chamber were stained with DPAI and streptavidin-AF647 for fluorescence microscopic imaging.

**4. Detection of iDC–OT-I CD8$^+$ T cell interaction by FT functionalized dendritic cells (DC-FT)**

iDCs generated from CD45.1$^{+/+}$ C57BL/6J mice were resuspended in HBSS buffer (100 µL per 1 million cell) containing 20 mM $MgSO_4$, 3 mM HEPES and 0.5% FBS. The cells were then treated with GDP-Fuc-FT (concentration as indicated in specific experiments). After 20 minutes of incubation on ice, FT functionalized iDC were washed with PBS twice and ready for use. The iDC-FT were then resuspended in T cell medium and left untreated or primed with $OVA_{257-264}$ or LCMV $GP_{33-41}$ (100 nM, different peptides or concentrations were used in certain experiments) at 37 °C for 30 min. After washing twice, DC-FT were co-cultured with purified

CD8+ T cells or splenocytes from CD45.1⁻ OT-I mice as a ratio of 1:1 (different ratios were used in some experiments) on a 96-well plate (20000 cells per well). After incubation for 2 hours (or indicated times) at 37 °C, GDP-Fuc-biotin (50 μM) were added gently and cells were incubated at 37 °C for a further 30 min. Then, LacNAc (final concentration 5 mM) were added to quench the reaction. After washing twice with PBS, cell mixture was analyzed by flow cytometry.

#### 5. Detection of selective interaction of dendritic cells in the mixture of OT-I $CD8^+$ T cells and P14 $CD8^+$ T cells

iDCs generated from $CD45.1^-$ C57BL/6J mice were resuspended in HBSS buffer (100 μL per 1 million cell) containing 20 mM $MgSO_4$, 3 mM HEPES and 0.5% FBS. The cells were then treated with GDP-Fuc-FT (0.2 mg/mL). After 20 minutes of incubation on ice, the FT functionalized iDC (iDC-FT) were washed with PBS twice. Then, iDC-FT were divided into two groups, resuspended in T cell medium and primed with $OVA_{257-264}$ or LCMV $GP_{33-41}$ at 37 °C for 30 min. After washing twice, DC-FT were co-cultured with CD8+ T cells from $CD45.1^{+/-}$ OT-I splenocytes and $CD8^+$ T cells from $Thy1.1^{+/-}$ P14 splenocytes at the ratio of 1:1:1 on a 96-well plate (30000 cells per well). After 2 hours of incubation at 37 °C, GDP-Fuc-biotin (final concentration 50 μM) were added and cells for another 30 min. Then, LacNAc (final concentration 5 mM) were added to quench the reaction. After washing twice with PBS, cell mixture was analyzed by flow cytometry.

#### 6. Identification and enrichment of TSA reactive TILs via FucoID

*Tumor lysate preparation*

A small portion of tumor tissue was mechanically dissociated. After cell counting, the resulting tumor cell suspension (5 to 10 × $10^6$/mL) were subjected to three quick frozen-thrown cycles. After centrifuging at 2000 × g for 10 min, the supernatant was collected, aliquoted and stored at -80 °C for priming iDC.

*Enrichment of lymphocytes from tumor cell suspension*

TILs were isolated by mechanically dissociating tumor tissue prior to centrifugation on a discontinuous Percoll gradient (GE Healthcare). The resulting lymphocytes were washed with PBS and cultured in complete T cell medium containing 100 IU/mL for FucoID labelling.

*FucoID protocol for TSA reactive TILs labeling*

iDCs were primed with the indicated antigen or tumor lysates as described in general information section. Then, iDCs were labelled with GDP-Fuc-FT (0.2 mg/mL) to generate iDC-FT following the general procedure. Enriched tumor lymphocytes from a tumor of C57BL/6 mice were co-cultured with iDC-FT at the ratio of 10:1 in complete T cell medium plus 20 mM $MgSO_4$. After 2 hours of co-culturing, GDP-Fuc-Biotin (50 μM) were added and cultured for another 30 mins. At last, LacNAc (5 mM) was added for 20 mins to quench the reaction. (The workflow indicating specific congenic markers is summarized in Figure 4B and Figure 5A) Subsequently, cells were collected for flow cytometric analysis or cell sorting. In brief, cells were incubated with mouse Fc-blocker (BD Biosciences) for 30 mins at room temperature, then stained with anti-mCD8a-PE, anti-mCD45.1-PB, anti-mPD-1-FITC and streptavidin-APC. FITC Rat IgG2aκ FITC Isotype Ctrl was used as a PD-1 isotype control. The gating strategy was described as below. For B16-OVA tumor model, $PD\text{-}1^-$ population: $CD8a^+/CD45.1^{+/+}/PD\text{-}1^-/Biotin^-$; $PD\text{-}1^+Bio^-$ population: $CD8a^+/CD45.1^{+/+}/PD\text{-}1^+/Biotin^-$; $PD\text{-}1^+Bio^+$ population: $CD8a^+/CD45.1^{+/+}/PD\text{-}1^+/Biotin^+$. For B16, E0771 and MC38 tumor model, $PD\text{-}1^-$ population: $CD8a^+/CD45.1^-/PD\text{-}1^-/Biotin^-$; $PD\text{-}1^+Bio^-$ population: $CD8a^+/CD45.1^-/PD\text{-}1^+/Biotin^-$; PD-

1$^{+}$Bio$^{+}$ population: CD8a$^{+}$/CD45.1$^{-}$/PD-1$^{+}$/Biotin$^{+}$. In some experiments, all CD8$^{+}$ TILs expressing PD-1 were sorted as the subset of total PD-1$^{+}$. For analyzing the markers of each subsets, cells were divided into two groups after incubating with mouse Fc-blocker. One group of cells were stained with Ghost Dye Violet 510, anti-mCD8a-PB, anti-mPD-1-FITC, streptavidin-APC, anti-mCD39-PE/Cy7, anti-mCD103-PE. Another group of cells were stained with Ghost Dye Violet 510, anti-mCD8a-PE/Cy7, anti-mPD-1-FITC, streptavidin-percp5.5, anti-mCD137-APC, anti-mTIM-3-PE and anti-mTCF1-PB.

**7. *In vivo* re-stimulation of different subsets of TILs from B16-OVA tumor**

iDCs (CD45.1$^{-}$) were used to labelled TILs from male C57BL/6 mice (CD45.1$^{+/+}$) with the procedures shown above. Then, TILs were sorted into three subsets according to cell surface marker PD-1 and biotinylation: PD-1$^{-}$, PD-1$^{+}$Bio$^{-}$ and PD-1$^{+}$Bio$^{+}$. After cell sorting, three subsets of TILs (50000 TILs per mice) were injected to WT male C57BL/6 mice (CD45.2$^{+/+}$) by tail vein injection respectively. After the TILs were rested for 24 hours *in vivo*, the recipient mice were infected with listeria bacteria expression OVA antigen (LM-OVA). After another 7 days, blood was collected from each mouse and stained with anti-mCD45.1-FITC, anti-mCD3-PB, anti-mCD8a-PE and H-2Kb/OVA$_{257\text{-}264}$ MHC tetramer-APC for TCR clone expansion analysis. After another 30 days, splenocytes from each mouse were also stained with the same panel of antibodies for TCR clone expansion analysis. At last, the only left antigen specific CD8$^{+}$ T cells from the splenocytes from PD-1$^{+}$Bio$^{+}$ recipient mice were enriched by CD8$^{+}$ isolation kit and were transferred to WT male C57BL/6 mice (CD45.2$^{+/+}$, 3 x $10^{6}$ cells per mice) by tail vein injection and the mice were infected with LM-OVA on the next day. Their blood was stained with anti-mCD45.1-FITC, anti-mCD3-PB, anti-mCD8a-PE and H-2Kb/OVA$_{257\text{-}264}$ MHC tetramer-APC for flow cytometry on day 8. (The workflow is summarized in Figure 4D)

**8. Rapid expansion of TILs**

Sorted TILs were rested in complete T cell medium with 100 IU/mL rhIL-2 for 1 day. Then, TILs (6000 TILs per well) were co-cultured with feeder cells (50 Gy irriadiated BALB/c mouse splenocytes, 0.6 x $10^{6}$ cells per well) in 300 μL complete T cell medium supplied with 500 ng/mL anti-mCD3 and 1500 IU/mL rhIL-2 in a 48-well plate. From day 5, cells were counted and fresh T cell medium containing same components were added to maintain cell density at 0.5-1×$10^{6}$ cells/mL. In most experiments, TILs were collected at day 9 or day 10 for analysis or *in vitro*/*in vivo* studies.

**9. IFNγ EliSpot assay for TILs reactivity analysis**

EliSpot assay were performed using a mouse IFNγ EliSpot kit (ab64029, Abcam). DCs were pulsed with indicated antigens or tumor lysate as described above. Then, different subsets of TILs were co-cultured with DCs primed with the indicated antigens respectively (For TILs from B16 tumor: 2500 TILs/10000 DCs; For E0771 tumor: 3000 TILs/12000 DCs; For MC38 tumor: 2500 TILs/10000 DCs) in a pre-coated 96-well plate (included in the assay kit). After 20 hours of co-culturing, the results of IFNγ secretion were measured according to the kit protocol (https://www.abcam.com/mouse-interferon-gamma-elispot-kit-ab64029.html). The pictures of each well were taken using Zeiss KS EliSpot reader and spot numbered were counted by hand using image J (1.50i).

**10. *In vitro* tumor killing assays of TILs**

B16-OVA-luc, B16-luc, E0771-luc or MC38-luc cells (stably transduced with firefly luciferase) were seeded in 96-well plate (10000 cells per well). Expanded TILs were co-cultured with the corresponding cancer cells in complete T cell medium at indicated effector/target ratios for 20 hours. The detection reagent was directly added to the medium in each well according to the manufactory's manual (Bright-Glo, Promega). Plate reader ($Synergy^{H4}$, BioTek) were used to measure the luminescence of each well.

**11. TCRβ sequencing, mRNA sequencing and analysis**

Sorted TILs were cultured in complete T cell medium supplied with 100 IU/mL rhIL for 24 hours. RNA samples were then extracted from 10000 cells of each population using $Picopure^{TM}$ RNA isolation kit (Qiagen RNase-Free DNase set were used to digest trace DNA). Samples were kept at –80 °C for TCRβ and mRNA sequencing. TCRβ sequencing was performed and analyzed by iRepertoire, Inc (https://www.irepertoire.com). The library for RNA-seq was prepared using SMARTseq HT kit and sequencing was performed using NextSeq500 sequencing platform. The reads were trimmed for the adapter sequences using cutadapt 1.18 with Python 3.6.3 and the trimmed reads were mapped to the reference genome using the STAR aligner 2.5.2a. Gene abundance was estimated with python 2.7.11, and HTSeq 0.11.0. PCA and the differential gene expression analyses between different cell populations was performed using DESeq2 package in R. Significantly different threshold was Benjamini–Hochberg FDR (p.adjust) $< 0.05$ and $|\log_2(\text{foldchange})| \geq 0.6$. Gene ontology over-representation analysis was performed using clusterProfiler package in R. GSEA analysis was performed using GSEA 4.0.3 (Broad Institute).

**12. Generation of gene signatures from the literature**

For up-regulated exhaustion gene signature, we used the up-regulated gene profiles of exhausted $CD8^+$ T cells by chronic infection compared to naïve $CD8^+$ T cells, which was published in Wherry et al. (Wherry et al. 2007). For naïve/memory gene module and activation/dysfunction gene signatures, we used the gene modules reported in Singer et al. (Singer et al. 2016). For cell cycle gene signature, we used the cell cycle gene set reported in Li et al. (Li et al. 2019). For up and down-regulated tumor specific antigen gene signatures, we used the gene profiles shared by tumor-specific $CD8^+$ T cells from early (day 8) and late stage (day 30) tumors and excluding the gene profiles of exhausted $CD8^+$ T cells by chronic infection, reported in Schietinger et al. (see Figure 5B in Schietinger et al. 2016).

**13. Statistical analysis**

Statistical analyses were performed using GraphPad Prism software (version 7.0). Comparisons over groups were analyzed using two-way ANOVA tests followed by Sidak's multiple comparisons test, and comparisons of multiple samples at one group were analyzed using two tailed t-test or one-way ANOVA followed by Tukey's multiple comparisons test. For cell-based experiments, three biological replicates were performed. In all figures, ns, $P > 0.05$; *$P < 0.05$; **$P < 0.01$; ***$P < 0.001$; ****$P < 0.0001$.

#### Supplementary Figures

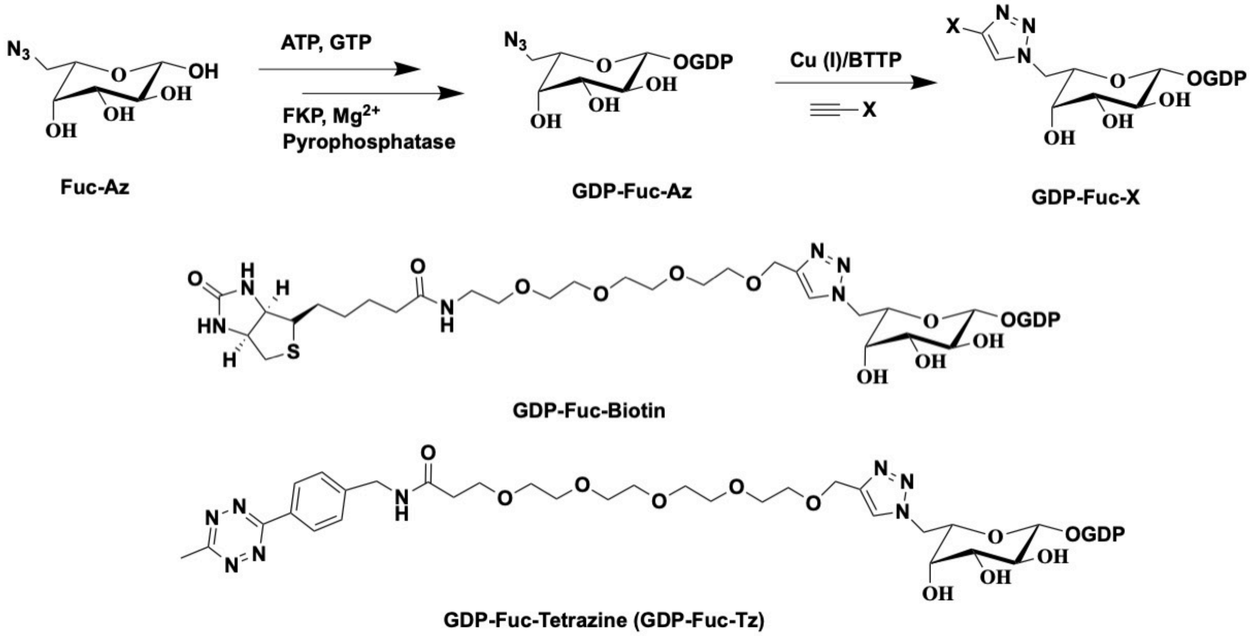


**Figure S1**. **Synthesis of GDP-Fuc-Biotin and GDP-Fuc-Tz**

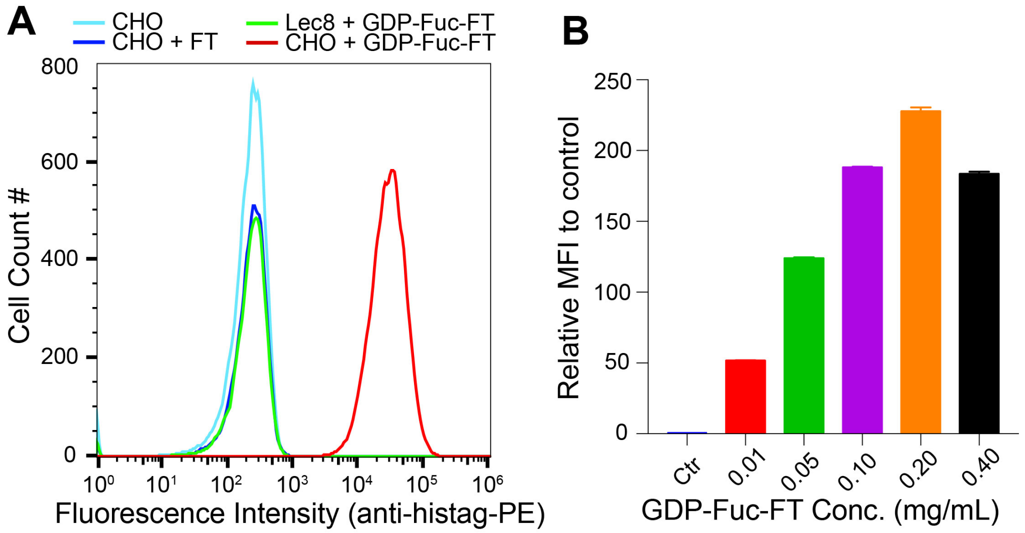


**Figure S2. Conjugation of the recombinant FT onto CHO cell surface.**
**(A)** Chinese hamster ovary (CHO) mutant Lec2 cells that express abundant LacNAc were treated with FT (0.1 mg/mL), GDP-Fuc-FT (0.1 mg/mL) or untreated (FT was expressed as a recombinant protein with a C terminal 6*His (histag)). In a control group, CHO Lec8 cells that do not express LacNAc were treated with GDP-Fuc-FT (0.1 mg/mL). Cells were incubated on ice for 20 min, followed by staining with DAPI and anti-histag-PE and analyzed by flow cytometry. Robust labelling was only achieved in Lec2 CHO cells treated with GDP-Fuc-FT. (**B**) Lec2 CHO cells were treated with GDP-Fuc-FT at different concentrations on ice for 20 min.

Then, cells were stained with anti-histag-PE for flow cytometry analysis. 0.20 mg/mL of GDP-Fuc-FT afforded the maximum FT labelling on CHO cells.

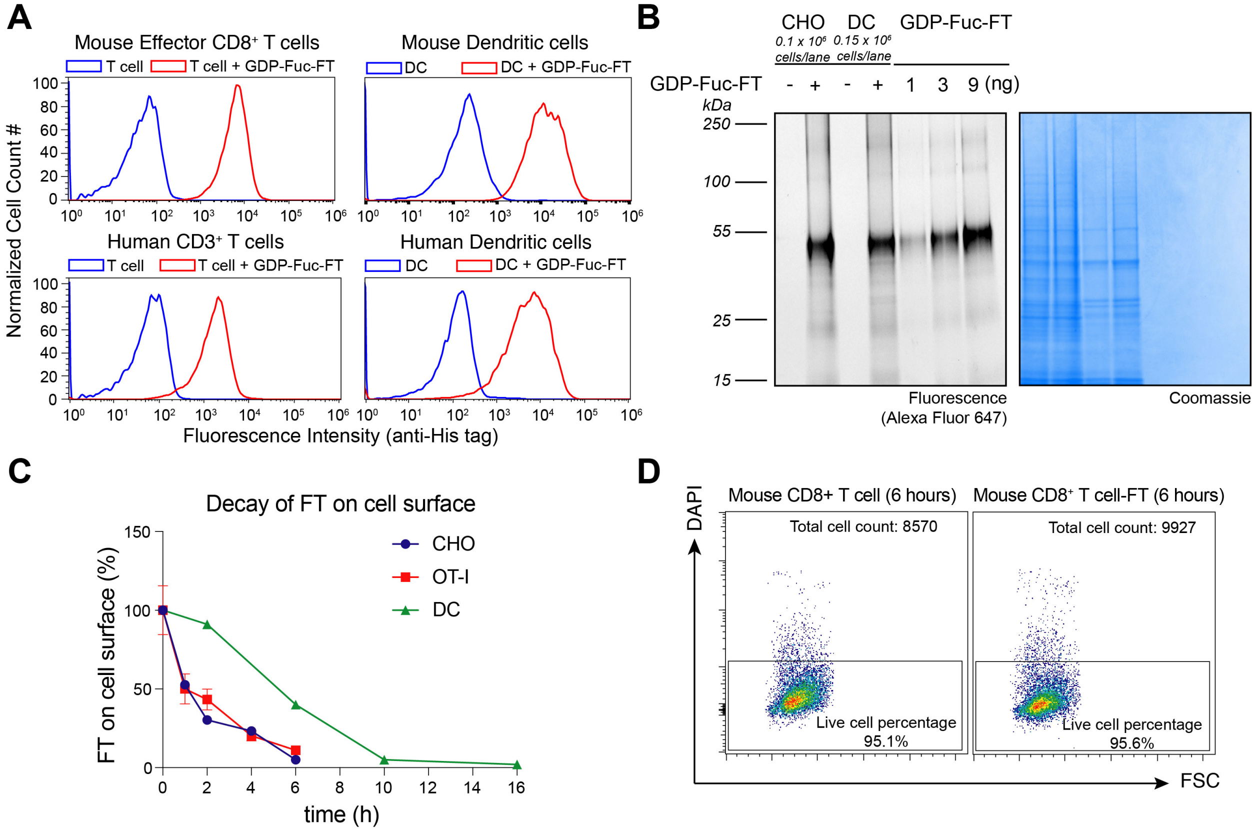


**Figure S3. Conjugation of FT onto multiple types of cells.**

**(A)** Mouse effector CD8$^+$ T cells from OT-I mice, mouse bone marrow dendritic cells (DCs) from C57BL/6 mice, human CD3$^+$ T cells and human DCs from PBMC were treated with GDP-Fuc-FT (0.2 mg/mL) for 20 min at room temperature or untreated. Then, cells were stained with anti-mCD8-PB (for mouse CD8+ T cells) or anti-mCD11c-APC (for mouse DCs) or anti-hCD8-FITC (for human CD3+ T cells) or anti-hCD11c-APC together with anti-histag-PE for flow cytometry analysis. **(B)** Fluorescent SDS-PAGE gel analysis of FT labelled cells. GDP-Fuc-FT modified with Alexa Fluor 647 (GDP-Fuc-FT-AF647) were used in the experiments for quantifying the number of FT molecules conjugated onto the cell surface. Different amounts of pure GDP-Fuc-FT-AF647 were used as standards in the quantification. At the GDP-Fuc-FT-AF647 concentration of 0.20 mg/mL, approximately 9 × $10^5$ FT molecules were introduced onto the surface of one CHO cell on average and approximately 5 × $10^5$ FT molecules were introduced onto one DC. **(C)** CHO cells, effector OT-I CD8$^+$ T cells and DCs were treated with 0.2 mg/mL GDP-Fuc-FT. Cells were stained with DAPI and anti-Histag-PE at different time post labelling for flow cytometric analysis of the decay of FT on cell surface. (**D**) 1 x $10^6$ Effector CD8$^+$ T cells generated from OT-I mice were treated with 0.2 mg/mL GDP-Fuc-FT for FT labelling. Same number of labelled cells and unlabelled cells were separately cultured in complete T cell medium containing 100 IU/mL rhIL-2 for 6 hours. Then, cells were stained with DAPI for flow cytometry analysis.

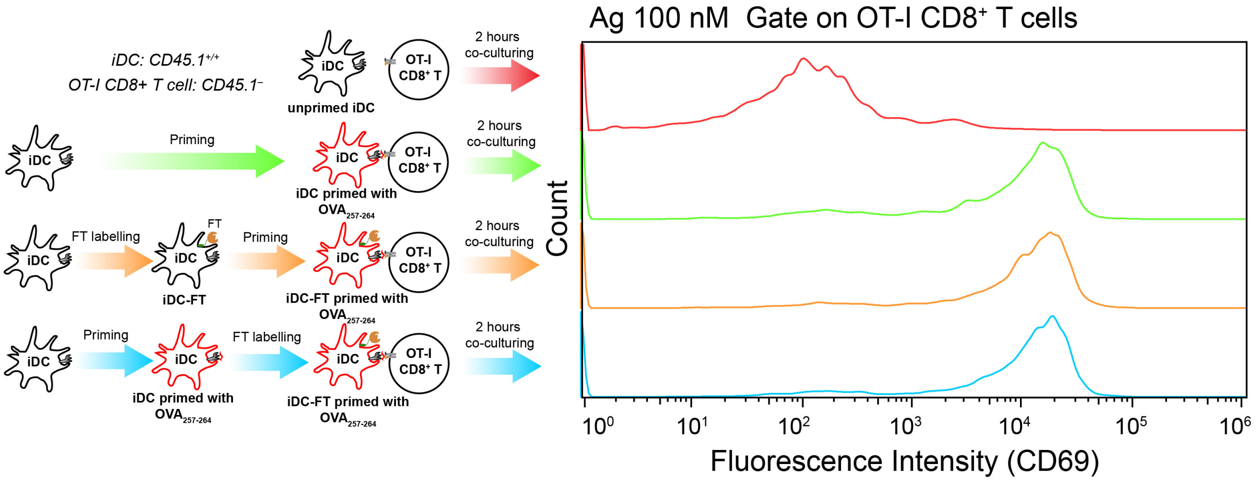


**Figure S4. Comparison of the antigen presentation abilities of iDC and iDC-FT**

iDCs derived from CD45.1$^{+/+}$ C57BL/6 mice were either labelled with FT then pulsed with OVA$_{257\text{-}264}$ or pulsed with OVA$_{257\text{-}264}$ then labelled with FT. They were then co-cultured with splenocytes from CD45.1$^{-}$ OT-I mice at cell ratio 1:1. Untreated iDCs loaded with OVA$_{257\text{-}264}$ or no antigen were used as a control. After 2 hours of co-culturing, cell mixtures were stained with anti-mCD45.1-FICT, anti-mCD8a-PB and anti-mCD69-PE for flow cytometry analysis, showing the FT functionalization of iDCs did not affect the iDC-mediated upregulation of CD69 in CD8$^{+}$ T cells. Representative flow cytometry figures from three replicates are shown.

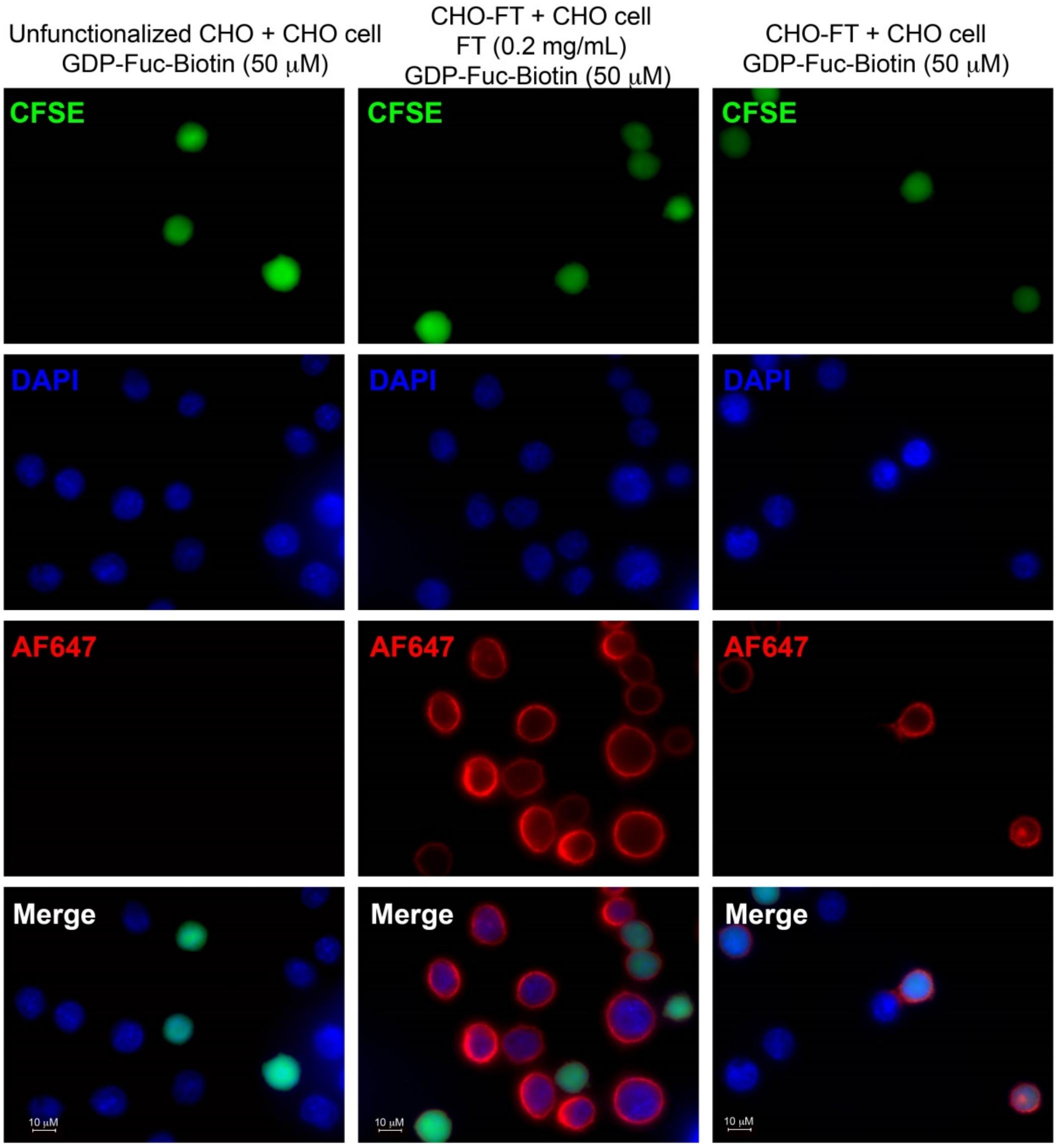


**Figure S5**. **Fluorescent imaging of FucoID mediated intercellular labeling of CHO cells.**
In the left column, untreated CHO cells (green) were co-cultured with CHO cells followed by adding GDP-Fuc-Biotin. No any biotinylation (red) was observed. In the middle column, CHO-FT (green) were co-cultured with CHO cells followed by the treatment of FT and GDP-Fuc-Biotin. Cell surface biotinylation (red) was observed on all cells. By contrast, in the right column, CHO-FT cells (green) were co-cultured with CHO cells followed by the addition of GDP-Fuc-Biotin. Fucosyl-biotinylation was observed on the interacting surface between CHO-FT and CHO cells. CHO cells without the contact of a CHO-FT were not biotinylated. In addition, CHO-FT exhibited robust self-biotinylation.

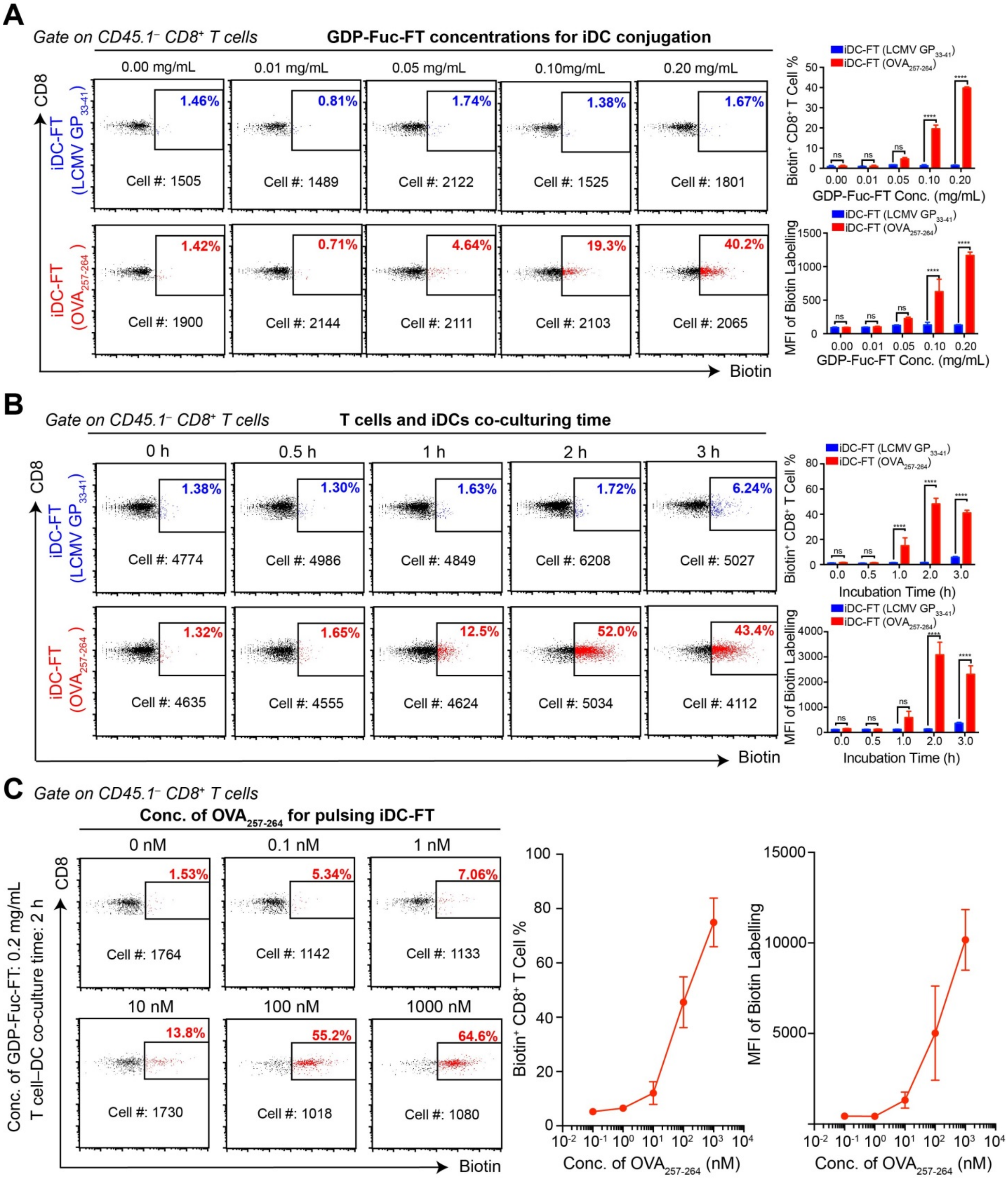


**Figure S6. Optimization of FucoID condition in the iDC-OT-I CD8⁺ T co-culturing system.** **(A)** Representative flow cytometric analysis of antigen specific fucosyl-biotinylation of OT-I CD8⁺ T cells by iDCs anchored with different amount of FT. iDCs ($CD45.1^{+/+}$) were treated with indicated amounts of GDP-Fuc-FT. Then, iDC-FT loaded with antigen $OVA_{257-264}$ or LCMV $GP_{33-41}$ (100 nM) were co-cultured with $CD45.1^{-}$ OT-I splenocytes at iDC/T ratio 1:1 for 2 hours followed the addition of GDP-Fuc-Biotin (50 μM) and another 30 minutes incubation. After quenching with LacNAc (2 mM), cells were washed and stained with anti-mCD45.1-FITC, anti-mCD8a-PB and streptavidin-APC for flow cytometry analysis. (**B**) Representative flow cytometric analysis of antigen specific biotinylation of OT-I CD8$^{+}$ T cells by iDC-FT with different co-culturing time before adding GDP-Fuc-biotin. iDCs (CD45.1$^{+/+}$) were treated with 0.2 mg/mL GDP-Fuc-FT. Then, iDC-FT loaded with antigen OVA$_{257\text{-}264}$ or LCMV GP$_{33\text{-}41}$ (100 nM) were co-cultured with CD45.1$^{-}$ OT-I splenocytes at iDC/T ratio 1:1 for the indicated time followed by the addition of GDP-Fuc-Biotin (50 μM) and another 30 minutes incubation. After quenching with LacNAc (2 mM), cells were washed and stained with anti-mCD45.1-FITC, anti-mCD8a-PB and streptavidin-APC for flow cytometry analysis. iDC/T ratio = 1:1. (**C**) Antigen dose dependent curve of the biotinylation of OT-I CD8$^{+}$ T cells by iDC-FT. Experiment procedure is shown in **Figure 3A** except iDC-FT were pulsed with indicated amounts of OVA$_{257\text{-}264}$. In all figures, mean ± SD (n=3); ns, P > 0.05; *P < 0.05; **P < 0.01; ***P < 0.001; ****P < 0.0001; two-way ANOVA followed by Sidak's multiple comparisons test.

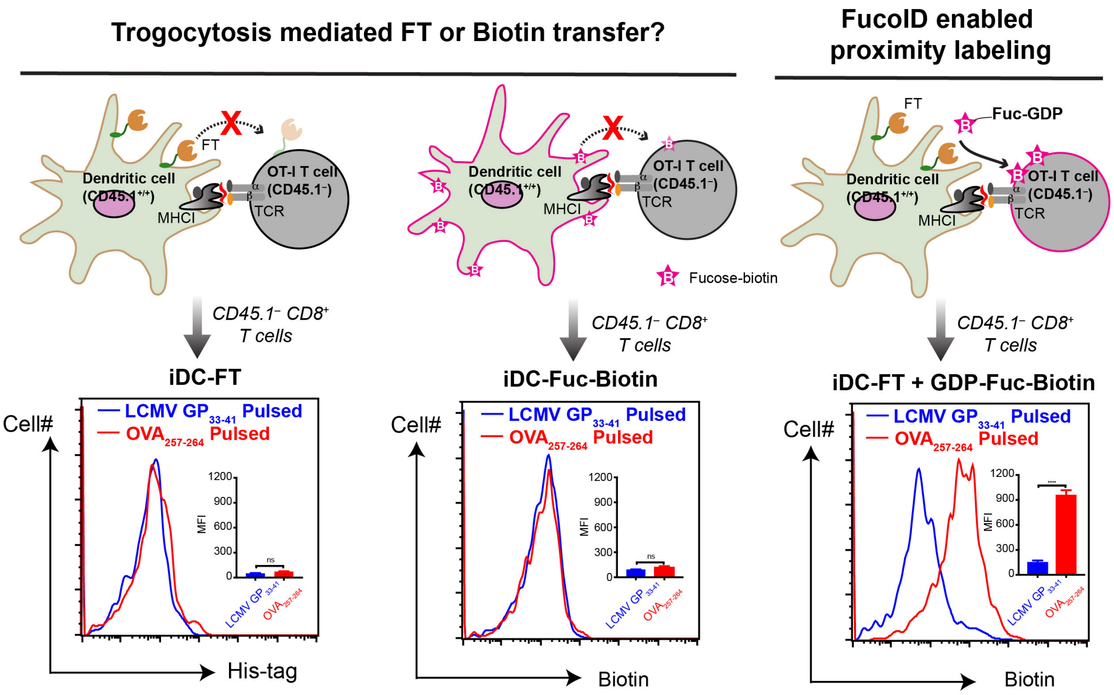


**Figure S7. Trogocytosis is not observed during antigen-dependent intercellular labeling via FucoID**

iDCs (CD45.1$^{+/+}$) were treated with 0.2 mg/mL GDP-Fuc-FT to generated iDC-FT or were treated with GDP-Fuc-Biotin (50 μM) and FT (0.2 mg/mL) to introduce biotin to iDC surface (DC-Fuc-Biotin). Then, iDC-FT pulsed with OVA$_{257\text{-}264}$ or LCMV$_{33\text{-}41}$ were co-cultured with OT-I splenocytes (CD45.1$^{-}$) to perform antigen dependent Fucosyl-biotinylation as the procedures in **Figure 3A**. Meanwhile, iDC-Fuc-Bio (prepared according to Li et al., 2018) pulsed with OVA$_{257\text{-}264}$ or LCMV$_{33\text{-}41}$ were co-cultured with OT-I splenocytes (CD45.1$^{-}$) for 2 hours. Cells were then stained with anti-mCD45.1-FITC, anti-mCD8a-PB, streptavidin-APC and anti-histage-PE for flow cytometry analysis. Mean ± SD (n=3); ns, P > 0.05; *P < 0.05; **P < 0.01; ***P < 0.001; ****P < 0.0001; two-tailed t-test.

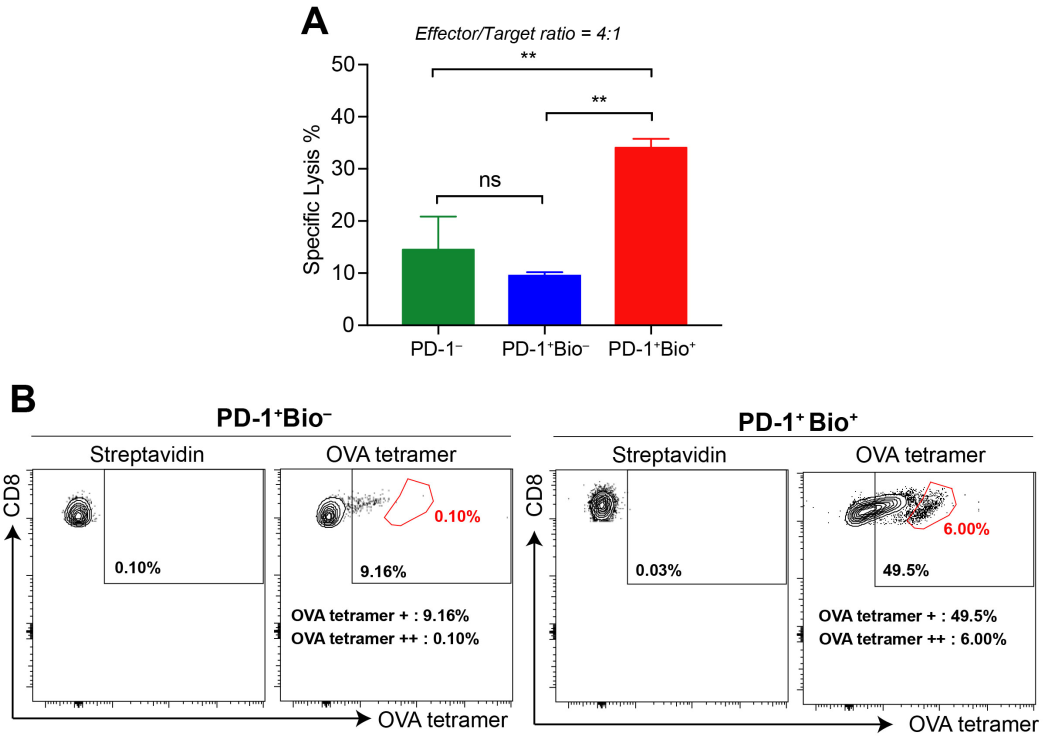


**Figure S8. Tumor reactivity and TCR specificity of PD-1$^+$Bio$^-$ and PD-1$^+$Bio$^+$ TILs isolated from murine B16-OVA tumor via FucoID, related to Figure 4.**

(**A**) Comparison of the expanded PD-1$^-$, PD-1$^+$Bio$^-$ and PD-1$^+$Bio$^+$ TILs in killing B16-OVA tumor cells at the effector-to-target ratio of 4:1. Labelling and sorting procedure are shown in **Figure 4B and C**. Sorted TILs from B16-OVA tumor were expanded under a rapid expansion protocol for 7 days before the analysis. Mean ± SD (error bars); n=3; ns, $P > 0.05$; *$P < 0.05$; **$P < 0.01$; ***$P < 0.001$; ****$P < 0.0001$; one-way ANOVA followed by Tukey's multiple comparisons test. (**B**) Comparison of the OVA specificity of PD-1$^+$Bio$^-$ TILs and PD-1$^+$Bio$^+$ TILs sorted from B16-OVA tumors. The isolated TILs were cultured in complete T cell medium supplied with 100 IU/mL rhIL-2 for 48 hours. The cells were then stained with anti-mCD8a-PE and H-2Kb/OVA MHC Tetramer-APC. TILs were stained with anti-mCD8a-PE and streptavidin-APC as a control to confirm biotin tag introduced via FucoID had disappeared on the cell surface.

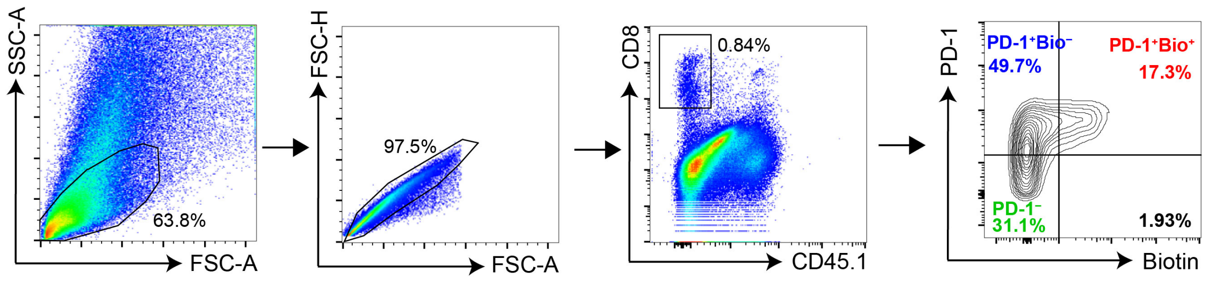


**Figure S9. Representative flow cytometry figures showing the gating strategy of E0771 TILs after FucoID labelling, related to Figure 5B.**

Similar Gating strategy were used for B16, E0771 and MC38 TILs.

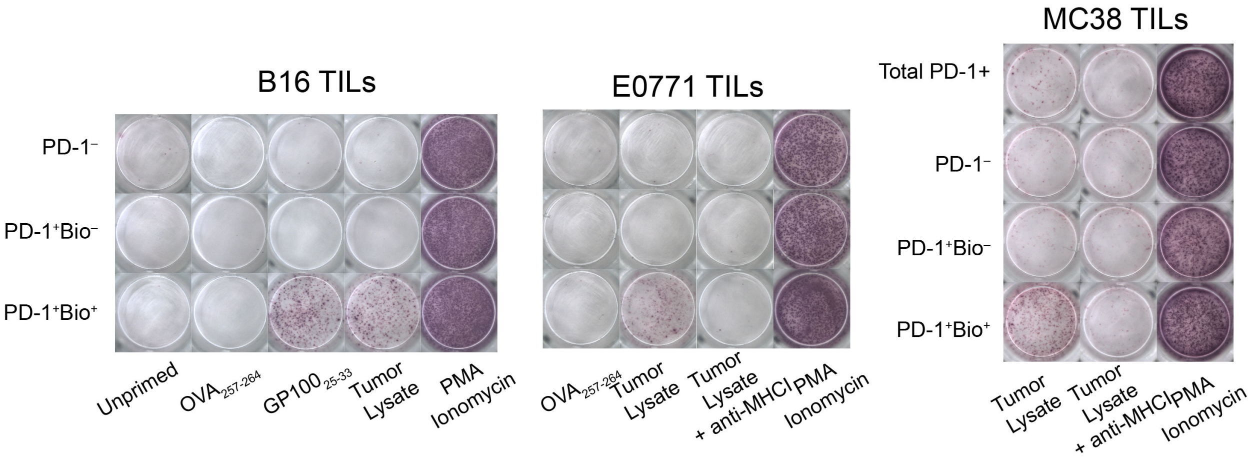


**Figure S10. Representative images of IFNγ EliSpots of TILs, related to Figure 5C.**

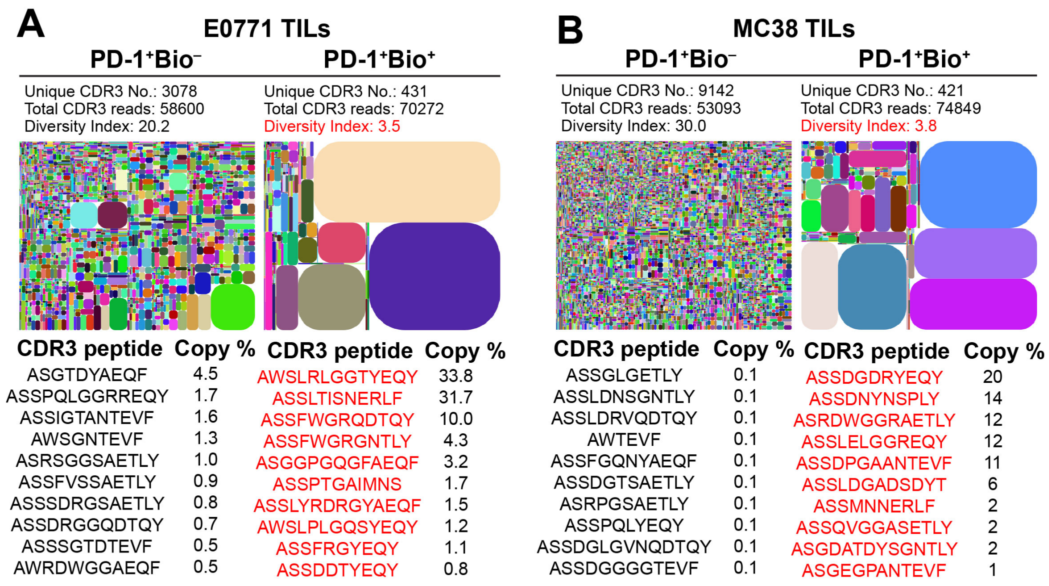


**Figure S11. Representative TCRβ RNA sequencing results of PD-1⁺Bio⁻ and PD-1⁺Bio⁺ TILs from E0771 tumor and MC38 tumor.**

Each spot in the plot represents a unique clonotype: V-J-CDR3, and the size of a spot denotes the relative frequency. The entire plot area is divided into sub-area according to V usage, which is subdivided according to J usage and then CDR3 frequency, subsequently. The top 10 most abundant CDR3 encoded peptide sequences in each population are shown. Unique sequences only found in PD-1⁺Bio⁺ population are highlighted in red. Three independent biological replicates were analyzed to confirm the TCR clonotype enrichment in PD-1⁺Bio⁺ TILs.

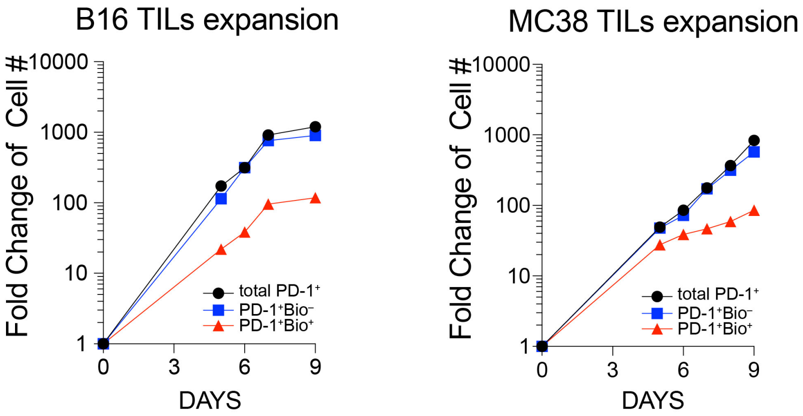


**Figure S12. Cell expansion curves of total PD-1$^+$, PD-1$^+$Bio$^-$ and PD-1$^+$Bio$^+$ TILs under the rapid expansion protocol.**

Representative expansion curves from three replicates are shown.

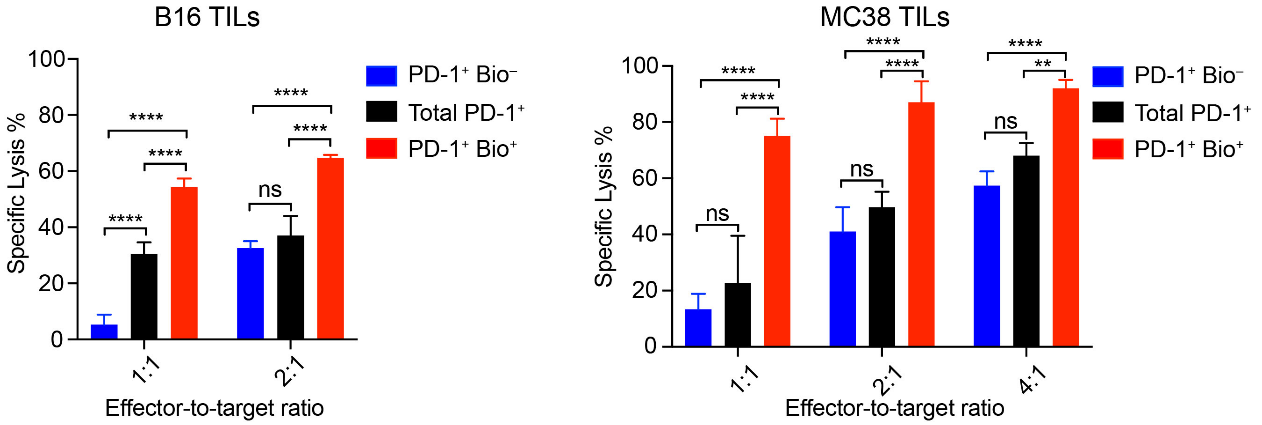


**Figure S13. Comparisons of the cancer cell killing activity of expanded total PD-1$^+$ TILs and PD-1$^+$Bio$^+$ TILs at different effector-to-target cell ratios, related to Figure 5D**.

Mean ± SD (error bars); n=3; ns, $P > 0.05$; *$P < 0.05$; **$P < 0.01$; ***$P < 0.001$; ****$P < 0.0001$; two-way ANOVA followed by Sidak's multiple comparisons test.

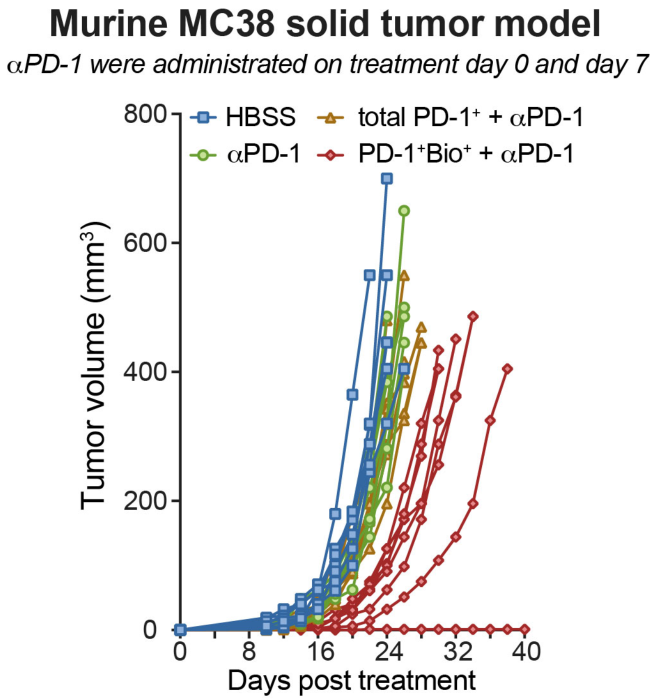


**Figure S14. Individual tumor growth of the MC38 model with adoptive TILs transfer treatment, related to Figure 5F.**

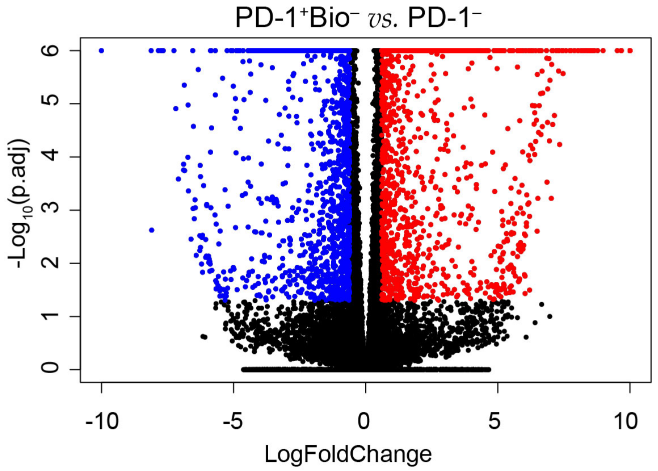


**Figure S15. Volcano plot of up- (red) and down- (blue) regulated genes of PD-1⁺Bio⁻ *vs.* PD-1⁻ TILs.**

Significance was determined as Benjamini – Hochberg FDR (p.adjust) $< 0.05$ and $|\log_2(\text{foldchange})| \geqslant 0.6$ ($\pm$ 1.5 fold).

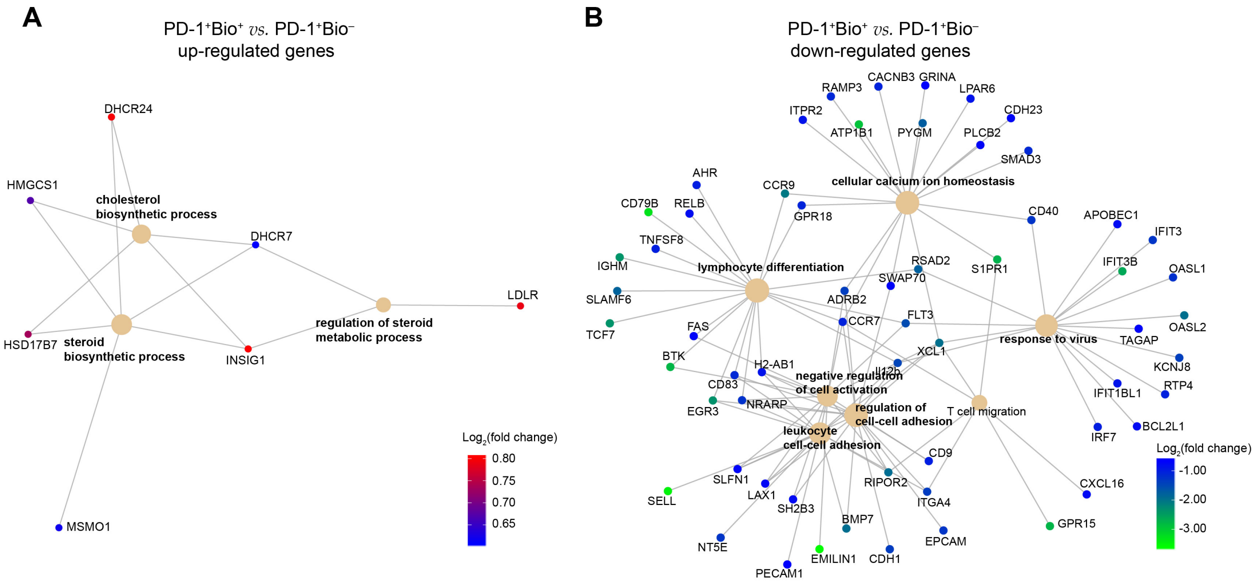


**Figure S16. Enriched biological process by up- and down-regulated genes in PD-1⁺Bio⁺ *vs.* PD-1⁺Bio⁻ TILs, related to Figure 6C.**

Gene concept networks was generated according to gene ontology (GO) over-representation analysis, showing genes involved in each enriched GO term.

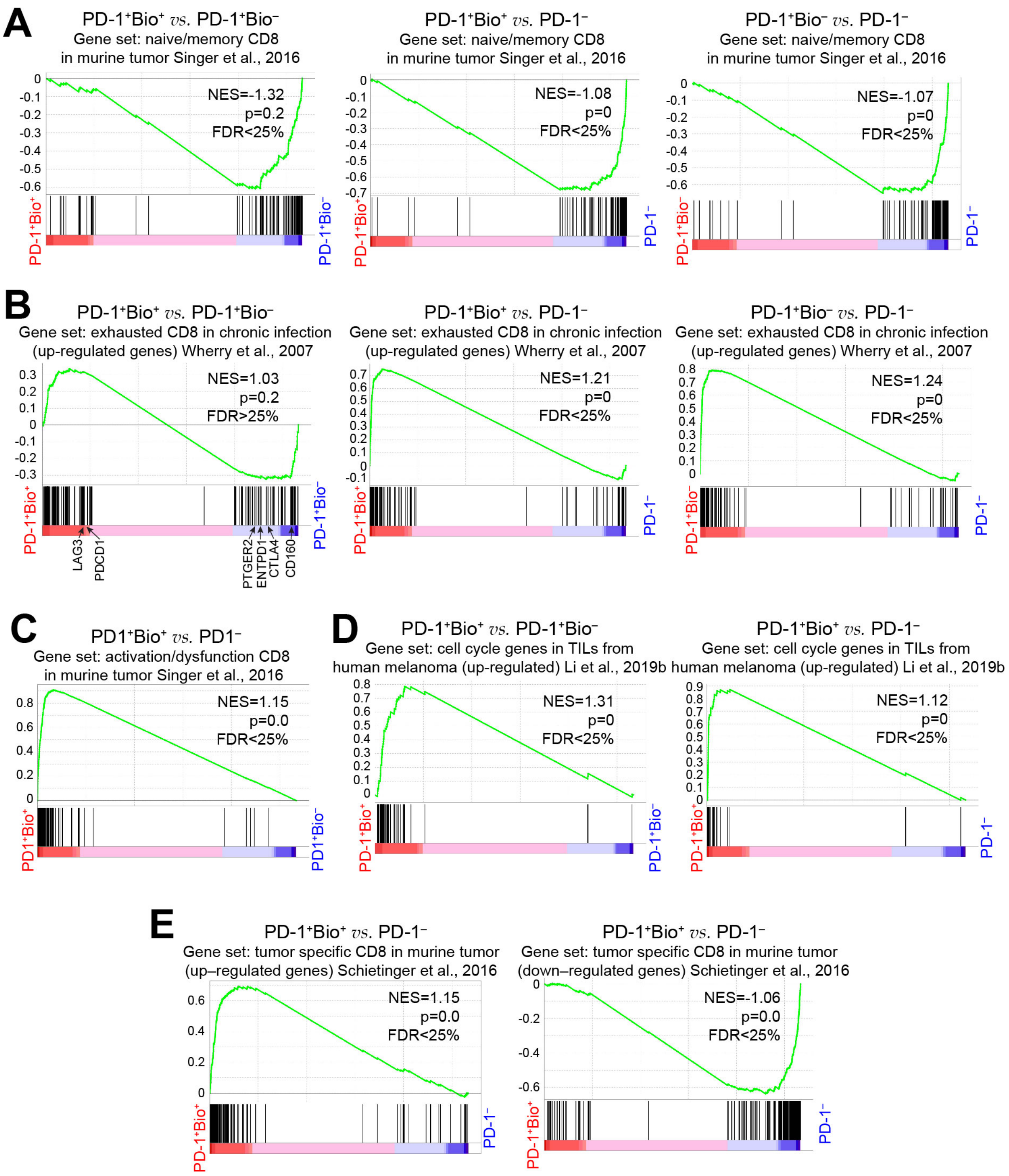


**Figure S17. Gene set enrichment analysis (GSEA) of the subsets of MC38 TILs using reported gene signatures, related to Figure 6D and E.**

(**A**) Three subsets were compared with each other, naïve/memory CD8 gene signatures were significantly enriched in the transcriptome of PD-1$^{-}$ TILs. (**B**) Three subsets were compared with each other. No enrichment of exhausted CD8 gene signatures were observed when comparing the transcriptome of PD-1$^{+}$Bio$^{+}$ with that of PD-1$^{+}$Bio$^{-}$ TILs. The transcriptome of PD-1$^{+}$Bio$^{+}$ and PD-1$^{+}$Bio$^{-}$ TILs showed similar enrichment of the exhausted CD8 signatures when comparing with that of PD-1⁻ TILs separately. (**C**) GSEA of activation/dysfunction CD8 gene module in the transcriptome of PD-1$^+$Bio$^+$ *vs.* that of PD-1$^-$ TILs. (**D**) GSEA of up-regulated cell cycle gene module in the transcriptome of PD-1$^+$Bio$^+$ *vs.* that of PD-1$^+$Bio$^-$ and in the transcriptome of PD-1$^+$Bio$^+$ vs. that of PD-1$^-$ TILs. (**E**) GSEA of up- and down-regulated tumor specific CD8 gene signature in the transcriptome of PD-1$^+$Bio$^+$ *vs.* that of PD-1$^-$ TILs.

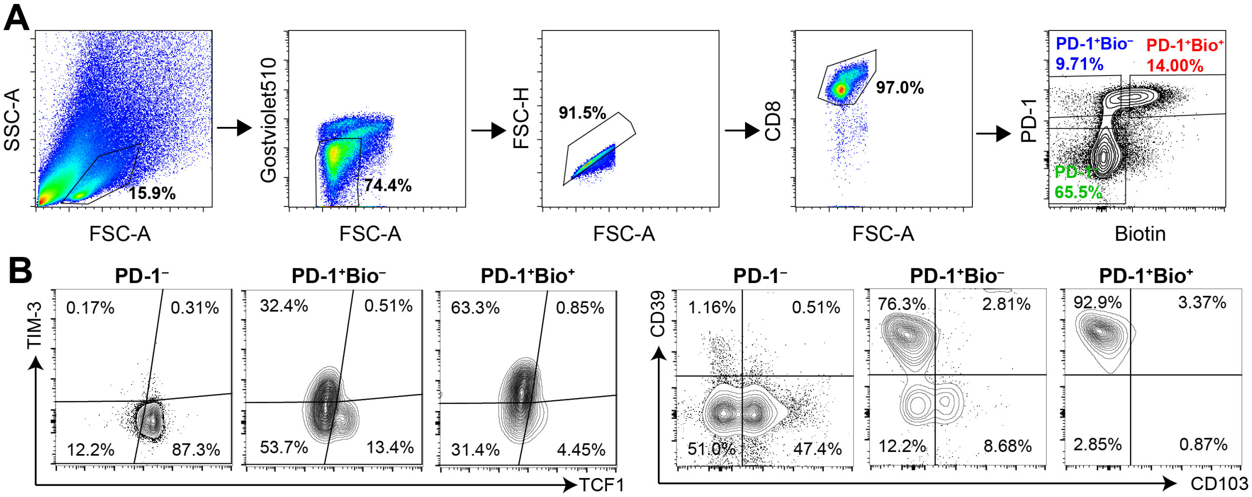


**Figure S18. Flow cytometric analysis of three subsets of CD8$^+$ TILs from murine MC38 tumor, related to Figure 6G.**

(**A**) Gating strategy for analyzing markers of PD-1$^+$Bio$^+$, PD-1$^+$Bio$^-$ and PD-1$^-$ TILs. (**B**) Representative flow cytometry analysis showing the staining of TIM-3, TCF1, CD39 and CD103 in the three subsets of TILs.
